# Distinct roles for motor cortical and thalamic inputs to striatum during motor learning and execution

**DOI:** 10.1101/825810

**Authors:** Steffen B. E. Wolff, Raymond Ko, Bence P. Ölveczky

## Abstract

The acquisition and execution of learned motor sequences are mediated by a distributed motor network, spanning cortical and subcortical brain areas. The sensorimotor striatum is an important cog in this network, yet how its two main inputs, from motor cortex and thalamus respectively, contribute to its role in motor learning and execution remains largely unknown. To address this, we trained rats in a task that produces highly stereotyped and idiosyncratic motor sequences. We found that motor cortical input to the sensorimotor striatum is critical for the learning process, but after the behaviors were consolidated, this corticostriatal pathway became dispensable. Functional silencing of striatal-projecting thalamic neurons, however, disrupted the execution of the learned motor sequences, causing rats to revert to behaviors produced early in learning and preventing them from re-learning the task. These results show that the sensorimotor striatum is a conduit through which motor cortical inputs can drive experience-dependent changes in subcortical motor circuits, likely at thalamostriatal synapses.

## Introduction

Whether tying our shoelaces or hitting a volleyball serve, we rely on our brain’s ability to learn and reliably generate stereotyped task-specific motor sequences. These processes depend on the interplay between cortical and subcortical brain areas, many of which have been identified (Fig. 1A)^1,2^. Yet, the specific roles of the different circuits and how they interact during motor sequence learning and execution is less well understood^1–3^. The striatum, the main input nucleus of the basal ganglia (BG), is an important nexus in the distributed mammalian motor control network, and one through which cortical and subcortical circuits can interact^4–8^. Its sensorimotor arm (dorsolateral striatum in rodents – DLS) receives excitatory input from sensorimotor cortex and thalamus. Striatal spiny projection neurons (SPNs), through their actions on BG output nuclei, can influence motor control at two levels (Fig. 1A). They can modulate motor cortical activity via the BG-thalamo-cortical pathway^4,9,10^, and further influence brainstem and midbrain motor circuits via direct projections from BG output nuclei (Fig. 1A)^4,11–17^.

**Figure 1:**
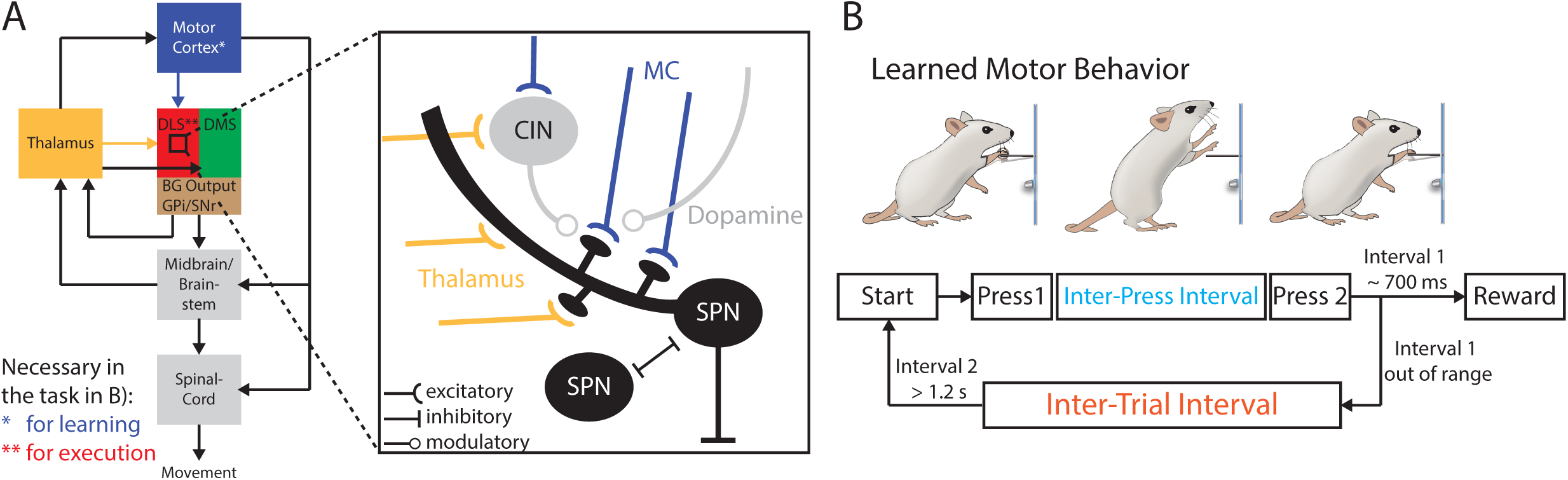
Probing the role of excitatory inputs to the striatum in motor sequence learning and execution. **A)** Simple schematic of the motor network probed in this study with the Basal Ganglia (BG) at its center. *Expanded view:* The main excitatory inputs to the Dorsolateral Striatum (DLS) are from Motor Cortex (MC, blue) and thalamus (yellow). Both inputs target overlapping populations of spiny projection neurons (SPNs) in the DLS, but also cholinergic (CIN) and other interneurons (not shown). **B)** Behavioral paradigm to test the role of the DLS and its excitatory inputs during the learning and execution of a complex motor sequence. Rats are rewarded for pressing a lever twice with a specific delay between presses (inter-press interval (IPI)). After unsuccessful trials animals must refrain from pressing the lever for 1.2 seconds (inter-trial interval (ITI)) to initiate a new trial.

While the involvement of the DLS in motor learning has been extensively probed^18–23^, its specific contributions, and especially how the inputs from motor cortex and thalamus enable these, remain poorly understood (though see companion paper^11^ and further references^24–30^). A common assumption, however, is that synaptic modifications at motor cortical inputs to the striatum play a role in memory storage^23,27,31–34^. Yet, the generality of this notion has been challenged by a recent study, showing that striatum is required for generating learned motor sequences that survive large bilateral motor cortex lesions^11^. Interestingly, the very same motor skills that can be executed without motor cortex, cannot be acquired without it^35^, suggesting a role for motor cortex in learning that is independent of its role in control. This raises the possibility that motor cortical inputs to striatum, rather than storing crucial aspects of the memory, guide learning-related plasticity in the subcortical motor network, possibly at non-motor cortex related synapses within the striatum^36^. Alternatively, motor cortex could exert its influence over the learning process by regulating plasticity in its other projection targets, such as the midbrain, brainstem and/or spinal circuits^37–47^.

If learning-related changes occur within the DLS but not at motor cortical inputs, it would suggest plasticity at DLS’s other main input – that from thalamus – as a possible mechanism for learning (Fig. 1A)^48–52^. DLS receives input mainly from the parafascicular nucleus (Pf) and the rostral intralaminar nuclei (rILN) of the thalamus^49,51–54^, nuclei which relay information predominantly from subcortical circuits^49,52,55^. While the inputs from thalamus to the striatum are comparable in strength^24,29,30^ and numbers to inputs from cortex^56,57^, surprisingly little is known about their function. The thalamostriatal projection has been implicated in modulating activity of cortical inputs to striatum via cholinergic interneurons (Fig. 1A), in the regulation of attention and behavioral flexibility, and in providing information to the BG about behavioral state and context for rapid behavioral adaptations^58–64^. Yet, whether they play a more direct role in motor learning and execution remains unclear.

The aim of this study is to elucidate how motor cortical and thalamic inputs to the striatum contribute to the acquisition and control of learned motor skills. To probe this, we take advantage of a paradigm that trains highly stereotyped and task-specific motor sequences in rats (Fig. 1B)^35^. As alluded to above, these motor skills, once learned and consolidated, are contingent on DLS but not on motor cortex^11,35^. Using a variety of targeted circuit manipulations, we show that the DLS and the motor cortical neurons that project to it, are required for learning the behaviors we train. We also find that synaptic changes in the DLS are an essential part of the underlying motor memory, and that thalamic neurons projecting to the DLS are necessary for executing the learned motor sequences. These results provide important new insights into how the interplay between motor cortex, thalamus and striatum underlies the acquisition and production of learned motor sequences.

## Results

### Learning of complex motor sequences requires the DLS, but not the DMS

The task we used in this study trains highly stereotyped motor sequences, that once learned are stably expressed over long periods of time^35^. We know that motor cortex is essential for learning the skills that emerge in our task, but not for executing them once acquired^35^. We recently showed that DLS is an essential part of the subcortical motor network that generates the learned skill^11^. However, whether DLS is obligatory for the learning process remains an open question. For example, removal of DLS prior to learning could prompt other parts of the motor system, including cortex itself, to ‘take over’ and cover for the DLS. Several studies, employing different learning paradigms than ours, have also suggested that the dorsomedial striatum (DMS), and its cortical input from prefrontal regions^5,6,65^, are essential for early learning^23,66–70^. The idea is that control is gradually ‘transferred’ from DMS to DLS as the animal’s behavior becomes increasingly ‘automatized’^18,20,22,23,27,33,68,70–73^.

To directly probe the contributions of DMS and DLS to learning the motor sequences acquired in our task, we performed excitotoxic lesions of either region in naïve animals (Fig. 2A, S1). The lesioned rats were then placed in our automated training system^74^ and trained using our standard protocol, which rewards animals with a drop of water for pressing a lever twice separated by a specific time interval (inter-press interval (IPI); target: 700 ms) (Fig. 1B)^35^. We compared the performance of lesioned animals to a cohort receiving DLS-targeted control injections (retrobeads or GFP-expressing AAVs; Methods). Control animals learned, over weeks of training, to produce IPIs around the prescribed 700 ms target with increasing precision (Fig 2B-C, S2B). Consistent with prior studies, the control animals ‘solved’ the task by developing complex, idiosyncratic, and spatiotemporally precise motor sequences (Fig. 2B-D)^11,35^. After unsuccessful trials, they additionally learned to withhold lever-pressing for 1.2 s (inter-trial interval or ITI), a requirement for initiating a new trial (Methods, Figs. 1B, 2B-C, S2B)^35^. All control animals reached our performance criteria for mastering the task in less than 30,000 trials (mean IPI within +/− 10% of target (700 ms) and CV of the IPI distribution below 0.25, Figs. 2D, S2B). DMS-lesioned animals reached these criteria after a similar number of training trials as control animals (19.494 +/− 10.208 (DMS) vs. 18.620 +/− 8.9141 (control) trials (mean +/− SD); Figs. 2B-D, S2B). We note that the performances of DMS-lesioned animals were slightly worse in the initial phases of learning, consistent with previous reports about a role of the DMS in early phases of rotarod learning^23,75^. Nevertheless, DMS-lesioned animals reached criterion at the same time as control animals, suggesting that the DMS is not essential for learning our task.

**Figure 2:**
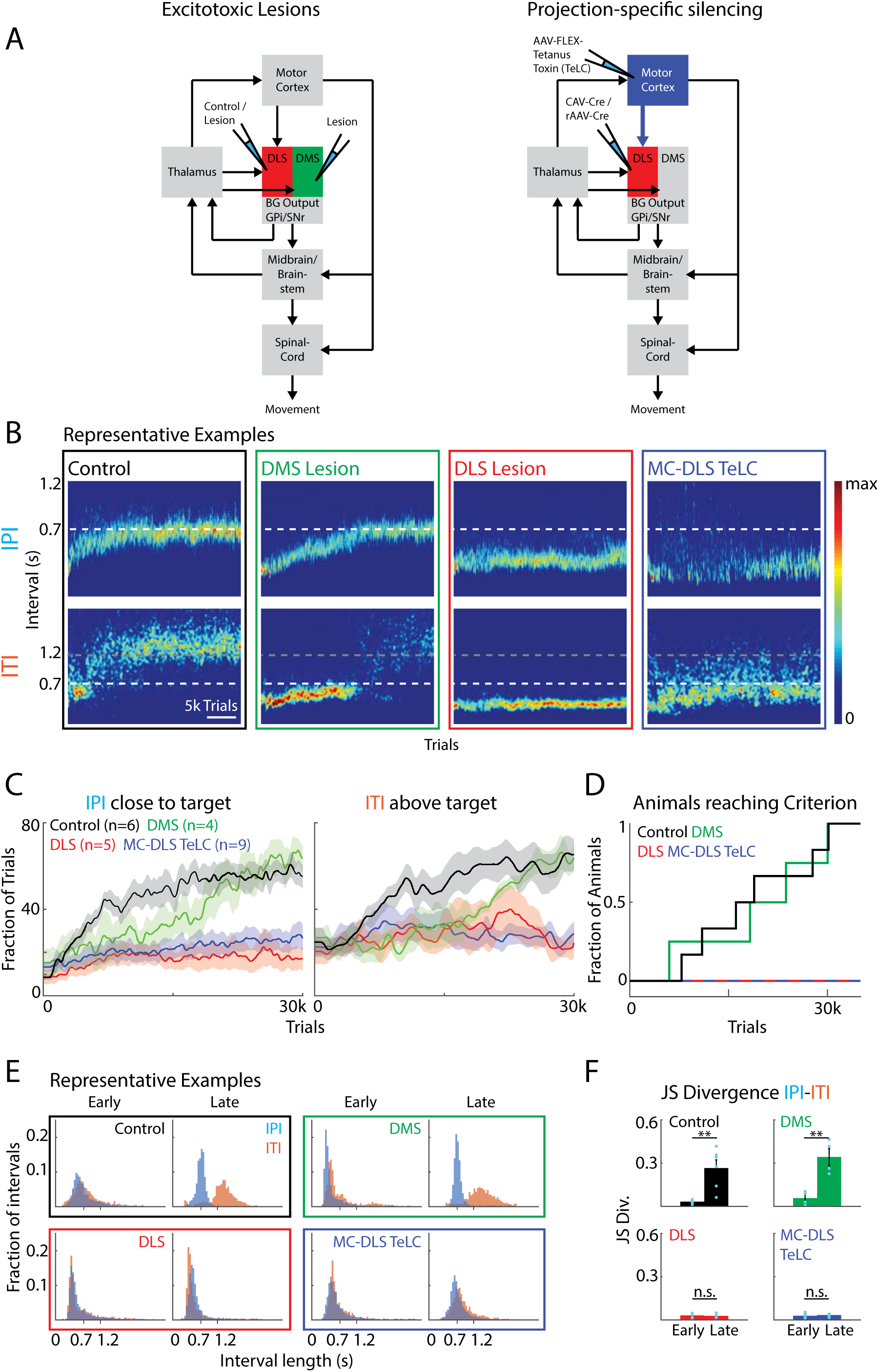
The DLS and its motor cortical input are required for learning complex motor sequences. **A)** Experimental schemes. *Left:* Excitotoxic lesions of either DLS or DMS, or DLS control injections in naïve animals. *Right*: Using a two-component viral strategy, we express Tetanus Toxin Light chain (TeLC) in DLS-projecting motor cortex neurons in naïve animals. Retrogradely infecting viruses for Cre expression are injected into the DLS and conditional AAVs for Cre-dependent expression of TeLC are injected into motor cortex. **B)** Representative examples of performance over the course of learning in animals subjected to different manipulations before the start of training (DLS control injection, DMS lesion, DLS lesion, silencing of motor cortex cells projecting to the DLS (MC-DLS) using Tetanus Toxin (TeLC)). Shown are heatmaps of the probability distributions of the IPIs and ITIs for representative example animals throughout learning. Colors indicate the probability of the occurrence of a certain interval in a given time windows (Methods). **C)** Averaged performance across animals for manipulations as in (B). *Left*: Fraction of trials with IPIs close to the target (700 ms +/− 20%). *Right*: Fraction of trials with ITIs above the threshold of 1.2 s. **D)** Fraction of animals reaching the learning criterion (see Methods) over the course of training. **E)** Distributions of durations between lever presses for the animals shown in (B) early (first 2000 trials) and late (trials 30,000 to 32,000) in training. **F)** Dissimilarity between the IPI and ITI distributions early and late in training as measured by the JS Divergence. Error bars represent standard error of the mean (SEM). \*\**P *< 0.01.

In stark contrast, none of the DLS-lesioned animals reached criterion performance in our task after 30,000 trials (Fig. 2B-D). A subset of animals for which training was extended for more than 60,000 trials also failed to reach our pre-specified criteria (Fig. S2C). Furthermore, while both control and DMS-lesioned animals learned the distinction between the IPI and ITI (Fig. 2E,F), DLS-lesioned animals did not distinguish between these intervals as indicated by overlapping IPI and ITI distributions (Fig. 2E,F). Together, our results show that the DLS, but not the DMS, is necessary for learning the task-specific motor sequences required to master our task and/or the strategy to deal with the underlying task structure.

### Learning requires motor cortex input to the DLS

Our finding that DLS-lesioned animals have learning deficits similar to animals with motor cortex lesions^35^, suggests the possibility that motor cortex informs learning-related plasticity in subcortical control circuits through its projections to the DLS. Motor cortical projections to control circuits in the midbrain and brainstem, which feed back to the DLS through the thalamus, provide a plausible alternative (Fig 1A)^12,14,16,40,44,76,77^. To more directly address the role of corticostriatal projections in learning, we used an intersectional viral approach (Fig. 2A)^78^ to silence DLS-projecting motor cortex neurons in naïve (untrained) animals. We injected retrogradely transported viruses expressing Cre-recombinase – either canine adenovirus type 2 (CAV-2)^79^ or retrogradely transported AAV (rAAV)^80^ – into the DLS. In addition, we injected a locally infecting adeno-associated viral vector (AAV), conditionally co-expressing Tetanus Toxin Light Chain (TeLC)^81–83^ and GFP in a Cre-dependent manner into motor cortex. TeLC cleaves the synaptic protein VAMP2 and thereby prevents the fusion of synaptic vesicles to the membrane and the release of neurotransmitters, effectively silencing neuronal output without killing the cells^81,84^. Our two-component viral approach restricts the expression of TeLC to motor cortex neurons with axon terminals in the DLS. By comparing the number of infected neurons in motor cortex with the number of neurons labeled by retrobeads co-injected into the DLS, we estimated our silencing rate to be about 50% of all DLS-projecting neurons in the infected parts of motor cortex (Methods). We used this approach to silence neurons in motor cortex that project to the DLS in a cohort of naïve animals (MC-DLS TeLC). Similar to DLS-(Fig. 2B-D) and motor cortex-lesioned animals^35^, MC-DLS TeLC animals failed to learn the task (Fig. 2B-D). They did not reach the learning criteria for the IPI within 30,000 trials (Figs. 2D, S2B), with a subset of animals undergoing extended training and not reaching the criteria after 60,000 trials (Fig. S2C). MC-DLS animals further failed to develop a distinction between IPI and ITI intervals (Fig. 2D-F). Together, these results show that motor cortex-originating corticostriatal neurons are necessary for learning the task-specific motor sequences we train, consistent with motor cortex guiding, or otherwise enabling, plasticity in subcortical motor circuits through direct projections to the DLS.

### Reversal of long-term synaptic plasticity in DLS disrupts performance

Once animals learn the stereotyped motor sequences required for our task, they are expressed in largely unaltered form over long periods of time^35^, suggesting the formation of stable memories. While prior studies have suggested that motor memories are stored in motor cortex^85–95^ and/or at synapses between motor cortex and striatum^23,27,31,32,34^, neither of these neural substrates are required for executing the behaviors we train^11,35^. The crucial involvement of the DLS in both motor sequence learning (Fig. 2) and execution^11^ suggest the possibility that the memory is stored, in part at least, at synapses in the DLS distinct from those formed by motor cortical inputs.

To address this, we took advantage of a pharmacological approach that reverses plastic changes at recently potentiated excitatory synapses, effectively erasing locally stored memory traces^96–101^. This approach relies on inhibiting the enzyme PKMzeta, which is crucial for maintaining synaptic potentiation^98,99,101^. Injection of its artificial inhibitor, ZIP, leads to synaptic depotentiation and degradation of learned behaviors contingent on plasticity in the targeted structure^33,96–99,102^. Skilled reaching, for example, is a dexterous and motor cortex-dependent behavior that is ‘unlearned’ after ZIP is injected into motor cortex^85,88^. Using this approach, we can pinpoint circuit elements that undergo task-relevant synaptic plasticity during learning and thus store some aspect of the memory necessary for executing the acquired motor sequences. To test whether the DLS is indeed a site of task-specific memory formation, we injected ZIP into this structure in expert animals (Figs. 3A, S3). We compared the outcome with ZIP injections into either motor cortex or the DMS (Figs. 3A, S3), neither of which is required for executing the learned motor sequences we train in our task^11,35^ and therefore not expected to store critical memories.

**Figure 3:**
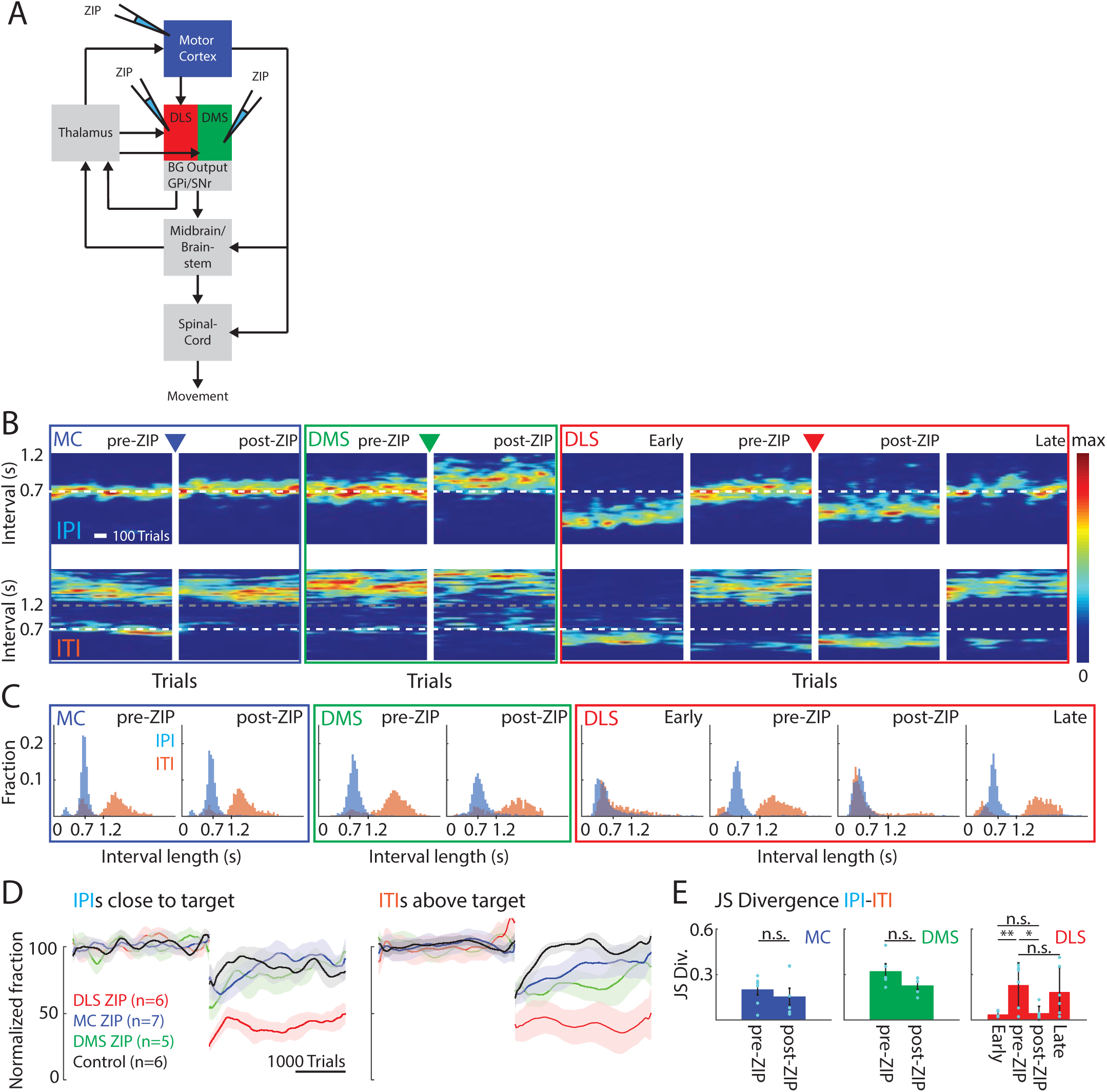
Pharmacological reversal of plasticity at excitatory synapses in DLS disrupts the execution of the learned behavior, but the same manipulation has no effect in motor cortex or DMS. **A)** Experimental scheme. ZIP, an inhibitor of an enzyme necessary for maintaining synaptic plasticity at excitatory synapses, was injected in either motor cortex, DMS or DLS. **B)** Representative examples of expert animals injected with ZIP in either motor cortex, DMS or DLS. Shown are heatmaps of the probability distributions of IPIs and ITIs before and after-ZIP for motor cortex and DMS injections. Additionally, performance early in training (first 2000 trials) and late (10,000 trials) after ZIP injection is shown for DLS ZIP animals. **C)** Distributions of durations between lever presses for the animals shown in (B) before and after ZIP injections. Early and late stages are additionally shown for the DLS ZIP animal. **D)** Population results for manipulations as in (B), normalized to performance before the manipulation. *Left*: Fraction of trials with IPI close to the target (700 ms +/− 20%). *Right*: Fraction of trials with ITI above the threshold of 1.2 s **E)** JS Divergence as a measure of dissimilarity between the IPI and ITI distributions, before and after ZIP injections. The reduction of the JS Divergence after DLS ZIP indicates an increase in the overlap of the two interval distributions (IPI and ITI) to levels observed early in training. *P < 0.05, **P < 0.01.

As expected, ZIP injections into motor cortex and the DMS did not affect task performance of expert animals beyond transient dips related to the effects of surgery (Figs. 3B,D, S4B). Performance measures and IPI and ITI distributions were similar to pre-ZIP training sessions in both cases (Figs. 3B-E, S4B). In contrast, ZIP injections into the DLS disrupted performance (Figs. 3B-E, S4B), despite the inhibitor only reaching parts of the DLS (Methods, Fig. S3). The distinction between IPI and ITI distributions was lost (Fig. 3C,E) and performance dropped to levels similar to early phases of training (Figs. 3B-C, S4B). Beyond ‘erasing’ recent memory traces, ZIP is not known to cause any permanent changes to circuit function^99,103,104^. To verify that the circuitry in the DLS was indeed intact after our injections, we re-trained injected animals on our task. After continued training DLS-ZIP animals reached pre-ZIP performance levels (Figs. 3B,C,E, S4B,C), indicating the formation of new task-specific memories.

These results suggest that at least part of the motor sequence memory is formed and stored in the DLS. While our approach does not reveal the exact identity of the potentiated synapses, it implicates excitatory synapses, since those are the ones affected by ZIP^98,99,105^.

### DLS-projecting thalamic neurons are necessary for task execution

These results alongside our previous study show that the DLS is necessary both for learning (see above, Fig. 2) and executing^11^ complex motor sequences and likely stores aspects of the motor memory (Fig. 3). But how does the DLS integrate into the larger motor network to serve these functions? A recent model of how cortical and thalamic inputs to striatum contribute to generating sequential behaviors suggested that learned sequences could be stored in intra-striatal synapses^106^. Because striatum is mostly an inhibitory structure, generating the requisite dynamics in its output neurons requires excitatory drive, which the DLS can receive from cortical, thalamic, and - for the direct pathway - nigral inputs^5,6,65^. Our ZIP experiments showed that excitatory inputs to DLS provide more than just a permissive excitatory drive. Indeed, our results suggested that they are a storage site of the motor memory. Since motor cortical input to DLS is dispensable for executing the learned behaviors^35^, these findings motivated us to look more closely at thalamic inputs to the DLS.

To probe the contribution of thalamic inputs, we silenced DLS-projecting thalamic neurons in expert animals. We targeted the main input nuclei to the DLS, the parafascicular, rostral intralaminar and midline nuclei^49,51–54^ (Figs. 4A, S5). We used the same intersectional viral strategy as before (Fig. 2), injecting retrogradely transported viruses for Cre expression into the DLS and AAVs for conditional TeLC expression into the thalamus (Figs. 4A, S5). As a control, we repeated the silencing of DLS-projecting motor cortical neurons (Fig. 2), but now in expert animals (Fig. 4A, S5). To further control for nonspecific effects of the surgery and viral expression, we expressed GFP in DLS-projecting neurons in motor cortex and thalamus in separate cohorts. Apart from transient effects due to the surgical procedure, performance in these control animals was not affected by GFP expression (Fig. 4B-E, S6).

**Figure 4:**
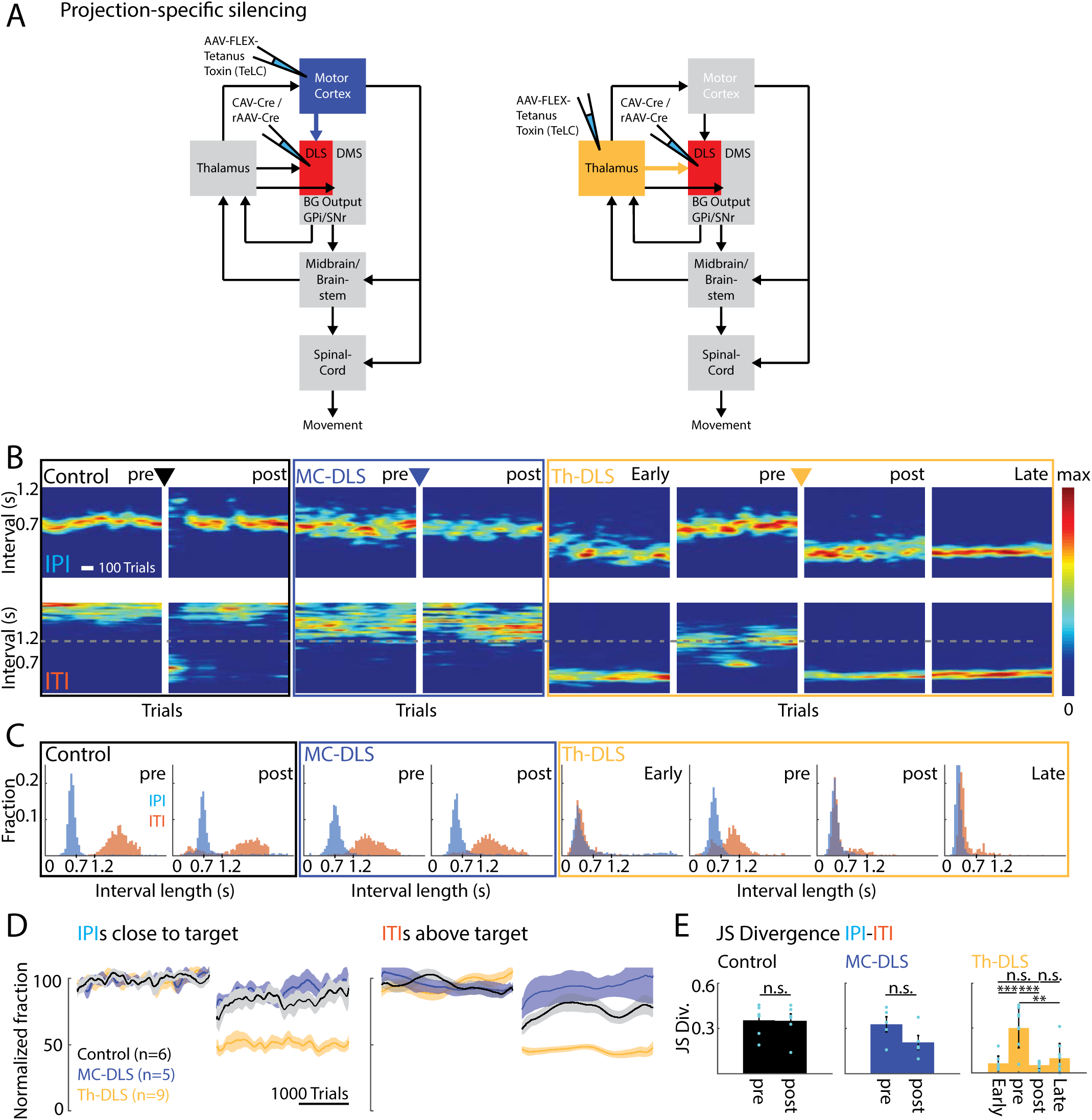
Silencing of DLS-projecting thalamus neurons in expert animals disrupts performance of the learned motor sequence, while silencing of the DLS-projecting motor cortex neurons has no effect. **A)** Experimental scheme. Using a two-component viral strategy, Tetanus Toxin Light chain (TeLC) was expressed in DLS-projecting neurons in motor cortex or thalamus. Retrogradely infecting viruses for Cre expression are injected into the DLS and conditional AAVs for Cre-dependent expression of TeLC are injected into either motor cortex or thalamus. **B)** Representative examples of expert animals which underwent silencing of neurons either in motor cortex (MC-DLS) or thalamus (Th-DLS) projecting to DLS or control virus injections. Shown are heatmaps of the probability distributions of IPIs and ITIs, pre- and post-silencing. **C)** Distributions of durations between lever presses for the animals shown in (B) before and after viral injections. **D)** Population results for manipulations as in (B), normalized to performance before the manipulation. *Left*: Fraction of trials with IPI close to the target (700 ms +/− 20%). *Right*: Fraction of trials with ITI above the threshold of 1.2 s. Controls include animals expressing GFP in neurons either in motor cortex or thalamus projecting to DLS (n=3/3). **E)** JS Divergence as a measure of dissimilarity between the IPI and ITI distributions, before and after viral injections. The reduction of the JS Divergence after Th-DLS silencing indicates an increase in the overlap of the two interval distributions to levels observed early in training. Shown are means and error bars represent standard error of the mean (SEM). **P < 0.01, ***P < 0.001.

In contrast to the learning effects we had seen (Fig. 2), but consistent with results from motor cortex lesions in expert animals^35^, silencing DLS-projecting motor cortex neurons did not impair task performance (Figs. 4B-E, S6). Intriguingly, however, silencing DLS-projecting thalamic neurons had very detrimental effects on performance (Figs. 4B-E, S6). While animals were still attentive to the task and pressed the levers during experimental sessions, they failed to produce the learned IPIs or ITIs (Figs. 4B,D, S6), and lost the distinction between the intervals altogether (Fig. 4C,E). Their post-silencing performance resembled early stages of training and did not recover even after extended periods of additional exposure to the task (Figs. 4B,C,E, S6).

To gain a better understanding of the role the thalamostriatal pathway plays in motor sequence execution, we precisely tracked the forelimb and head movements of a subset of animals using high-speed videography and automated markerless motion tracking (Methods)^107,108^. As described before^11,35^, animals developed highly stereotyped (Fig. 5A-D) and idiosyncratic (Fig. 5E-H) motor sequences over the course of training. Silencing the DLS-projecting thalamic neurons drastically altered the animals’ movements (Fig. 5A-D) and essentially phenocopied our previous excitotoxic lesions of the DLS (Fig. 5)^11^. Animals lost their learned stereotyped (Fig. 5A-D) and idiosyncratic (Fig. 5E-H) motor sequences, regressing to simpler lever pressing movement patterns that were similar across animals (Fig. 5E-H). Focusing on the lever-pressing movements themselves revealed that these were not only highly similar across animals (Fig. 5I,J) but also resembled movements used early in training (Fig. 5I,J), a behavioral reversion similar to what is seen after DLS lesions^11^. These results suggest that DLS-projecting thalamic neurons play an essential role in generating subcortically produced learned motor sequences.

**Figure 5:**
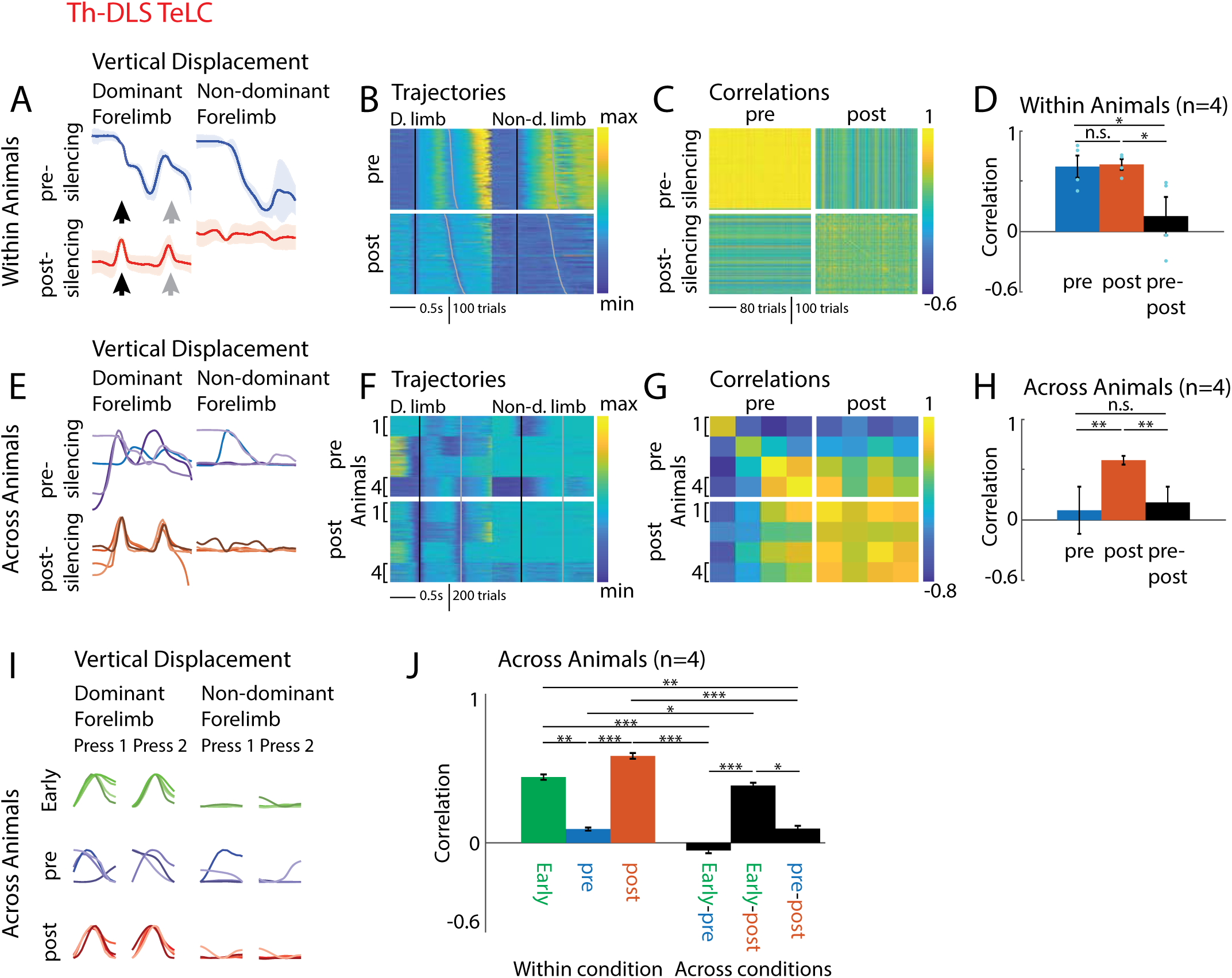
Silencing DLS-projecting thalamic neurons in expert animals leads to a loss of idiosyncratic learned motor sequences and a regression to simpler lever-pressing behaviors common across animals. **A)** Effects of Th-DLS silencing on lever-press related forelimb movements. Average forelimb trajectories (position in the vertical dimension) of a representative animal before (red) and after (blue) Th-DLS TeLC (calculated from trials close to the mean IPI (mean IPI +/− 30ms)). Black arrows indicate the 1^st^ lever press, grey arrows the 2^nd^ press. **B)** Vertical forelimb displacement in individual trials (color coded) before and after Th-DLS TeLC for dominant (forelimb used for 1^st^ lever press) and non-dominant forelimbs, sorted by IPI. Black lines mark the time of the 1^st^ lever press, grey lines of the 2^nd^ press. **C)** Pairwise correlations of the forelimb trajectories shown in B) after linear time-warping of the trajectories to a common time-base (see Methods). **D)** Average pairwise correlations within an animal averaged across animals. **E)** Comparison of average forelimb trajectories (vertical displacement) of all animals before and after Th-DLS silencing. **F)** Vertical forelimb displacement in randomly selected trials (150 per animal) of all animals before and after Th-DLS TeLC for dominant and non-dominant forelimbs, sorted by IPI. Black lines mark the time of the 1^st^ lever press, grey lines of the 2^nd^ press. **G)** Pairwise correlations between the trials shown in F), averaged per animal. **H)** Averages of the correlations shown in G) by condition (average of all pre-to-pre, post-to-post and pre-to-post correlations). **I)** Comparison of average forelimb trajectories (vertical displacement) of all animals early in training, before and after Th-DLS silencing. **J)** Averages of the correlations of all trials of all animals within and across the conditions in I). Shown for all bar graphs are Mean +/− SEM. *P < 0.05, **P < 0.01, ***P < 0.001.

## Discussion

Our study was designed to elucidate how thalamic and motor cortical inputs to striatum contribute to the acquisition and execution of task-specific motor sequences (Figs. 1, 6). We found that the DLS, but not the DMS, is required for learning, and that this function is contingent on motor cortical input to the DLS (Fig. 2). However, the very same input from motor cortex is dispensable for executing the learned motor sequences (Fig. 4), consistent with published results^35^. We identified plasticity at excitatory synapses in DLS as a likely substrate for the underlying motor memory (Fig. 3) and further showed that DLS-projecting thalamic neurons are essential for executing the consolidated behaviors (Figs. 4,5). These results are consistent with motor cortical inputs to DLS guiding plasticity at thalamo-striatal synapses, thus allowing subcortical motor circuits to learn and execute stereotyped task-specific motor sequences (Fig. 6). Taken together, these findings shed new light on the neural circuit-level logic by which motor skills are acquired, specifically the roles of DLS’s two major inputs, from motor cortex and thalamus (Fig. 6).

**Figure 6:**
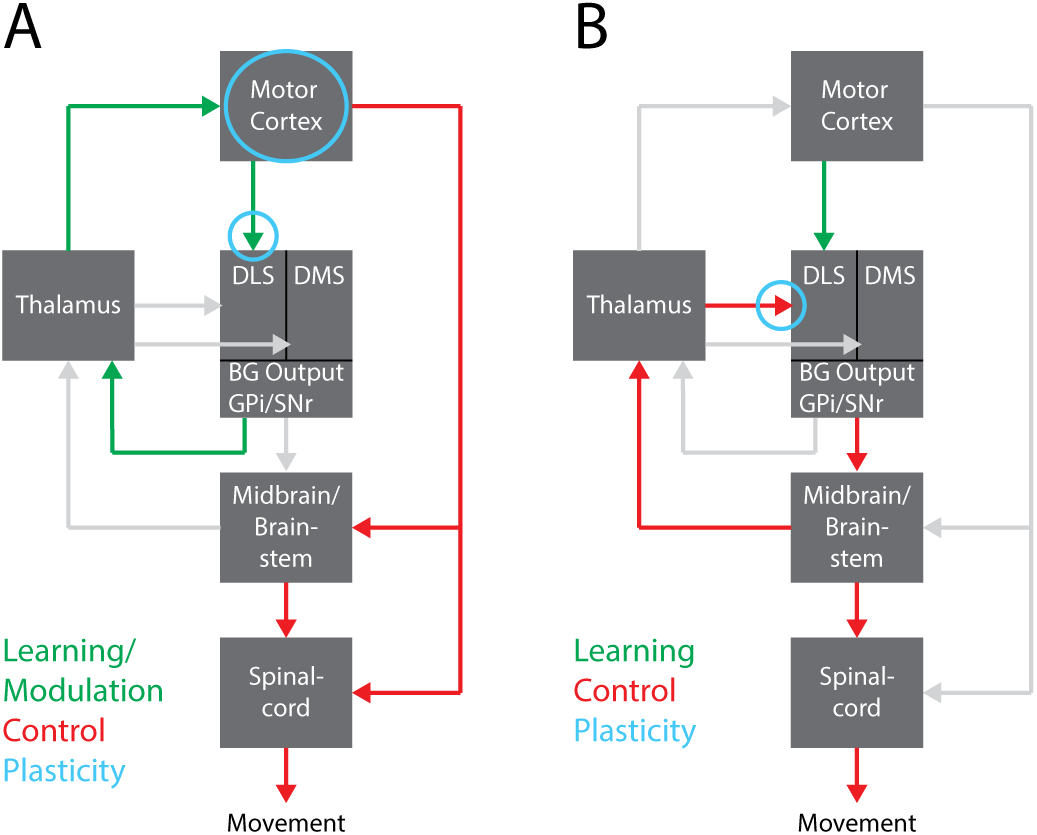
Alternative pathways for learning and execution of motor skills. **A)** Simplified view of the dominant circuit model for the learning and execution of motor skills. Motor Cortex plays a central role both for learning and control of motor skills. Motor cortex modulates the activity of the Basal Ganglia (BG) via its projection to the dorsolateral striatum (DLS). Over the course of learning this leads to plasticity at corticostriatal synapses. This allows the BG to modulate MC’s activity and output via the cortico-BG-thalamo-cortical loop in a more targeted manner. This, in turn, leads to plasticity within motor cortex, allowing motor cortex to control and effectively drive the execution of the desired, learned behaviors via its direct projections to motor control centers in the midbrain, brainstem and spinal cord. **B)** Our results suggest a different pathway for the learning and execution of motor skills. During learning, motor cortex ‘tutors’ the BG via its projections to the DLS. At least a part of this tutoring may be the gating or inducing of synaptic plasticity at thalamic input synapses to the DLS. Once the sequence has been learned, the subcortical circuitry, involving the BG-brainstem-thalamo-BG loop, is sufficient to drive the execution of the learned sequence and motor cortex input becomes dispensable. These pathways may interact to different degrees, depending on the behavior and the need for cortical involvement.

### Evolutionary considerations

Studies probing the corticostriatal pathway in motor learning have often assumed that motor cortex controls the acquired behavior^2,3,32,109–111^, making it difficult to distinguish its separate contributions to learning and control processes. In contrast, the motor sequences we train in our task have no explicit requirements for dexterity or cognition^112^, and have been shown to be motor cortex independent^35^. These highly stereotyped and task-specific behaviors, therefore, are likely stored and generated subcortically^11,35^. The involvement of both the BG^11^ and the thalamus in the execution of these learned behaviors (Figs. 4,5) implicates the phylogenetically older BG pathway that connects thalamus to BG to subcortical motor centers^7,12,14,113–116^.

It is widely assumed that this subcortical pathway is involved in initiating^1,12–14,114,117^ and modulating^118–122^ innate behaviors, such as locomotion, grooming, hunting and feeding^120,123–125^ through selective and graded disinhibition of its downstream targets. Our results suggest that the utility of this BG pathway can be extended to generate novel task-specific motor sequences - a process that may require a motor cortex-dependent reprogramming of subcortical motor circuits. This would allow cortex to off-load the ‘task’ of generating specialized and stereotyped learned behaviors to subcortical circuits, making them more automatized and less prone to cortical ‘interference’^126^. Consistent with this idea are observations that task-related cortical activity decreases over the course of training and with increasing automaticity^27,127–133^. Our current study suggests that the striatum may be where information from cortex is ‘handed-off’ to subcortical circuits.

The idea that one brain area ‘tutors’ another has been advanced also in other contexts^3,134–136^. Perhaps most relevant to our study is the example of vocal learning in songbirds, where a basal ganglia-related circuit guides plasticity in the connections between two premotor areas^137–139^. Although the circuit-level logic underlying the acquisition of stereotyped complex motor sequences in songbirds and rats is shared, we find that the functions of the homologous brain regions differ markedly. In songbirds, the role of the song-specialized BG mirrors that of motor cortex in our rats, whereas the BG (or DLS) in rats seem to have a control function similar to what has been attributed to the ‘cortical’ motor pathway in songbirds^140^. This could be yet another example of convergent evolution, in which a successful solution, in this case for learning complex stereotyped motor sequences, has evolved independently in different animals, resulting in similar principles and solutions but with different neural circuit elements^141^.

A mechanism for adapting ‘hard-wired’ species-typical behaviors to individual needs would certainly confer a fitness benefit and hence be favored by evolution. An evolving motor cortex may thus have been a boon to our ancestors not only because it enabled more sophisticated motor control strategies^2,77,91,142–145^, but also because it endowed a well-tuned and sophisticated subcortical motor control infrastructure with the capacity to meet new motor challenges.

While most studies, particularly in non-human primates and humans, focus on aspects of behavior in which cortical control is assumed to be essential, there is no reason to believe that the control of motor behaviors are either cortical or not. Rather, subcortical and cortical control circuits likely function in concert^77,142,145,146^. However, by studying a motor cortex-independent behavior, we can probe how the interplay between cortical and BG circuits adds functionality to subcortical motor circuits, and the mechanisms and pathways through which this is accomplished.

### Mechanisms for tutoring

Though we have identified the importance of motor cortical inputs to the striatum for learning the motor sequences we train in our task, the nature of this input and how it guides plasticity in striatum still remains to be understood. One possibility, analogous to the plasticity mechanism thought to underlie aspects of vocal learning in songbirds^139^, is that motor cortex guides plasticity at thalamostriatal synapses through spike-timing dependent heterosynaptic plasticity ^139,147,148^. Since the inputs from motor cortex and thalamus target overlapping populations of SPNs (Fig. 1A)^56,149^, precisely timed co-activation of these inputs could induce plasticity^147,148^. Such co-activation could be triggered by shared inputs to striatum-projecting motor cortex and thalamus neurons. Induction of plasticity at thalamic inputs may also require initial plasticity at cortical synapses – and both processes could be further modulated by dopamine^150–155^.

Another possibility for how motor cortical and thalamic input to striatum could work together to drive learning and retention of novel motor sequences is suggested by a recent modeling study^36^. In this work, cortical input to the DLS initially drives motor output and undergoes fast, dopamine-dependent plasticity^151,153–155^. Thalamic inputs to striatum do not drive behaviors at first but are increasingly strengthened via slow Hebbian plasticity at SPNs that are co-activated by the potentiated cortical input. Because of this continued strengthening, the thalamic inputs eventually assume control over the learned behavior, making motor cortex inputs redundant. Future studies will be necessary to probe the plasticity rules and mechanisms in striatum and how they underlie different aspects of motor learning^148,153,156^.

### Redundancy in motor networks

Though the redundancy between thalamic and cortical inputs for controlling striatal activity and behavior is an intriguing concept^36^, we see no evidence for it. Assuming, as is commonly done, that motor cortex is the main controller of learned motor skills, one could attempt to explain why the behaviors we train survive motor cortical lesions by suggesting that other circuits step in to ‘take over’ motor cortex’s control function^157^. Given that DLS is essential for control, this would require an equivalency between thalamic and cortical inputs to striatum in controlling the behavior. There isn’t one. Silencing thalamic input completely disrupts the learned behavior, while the same manipulation to cortical inputs has no effect (Figs. 4, S6). This suggests that the signal flow from thalamus to striatum, and likely plasticity at thalamostriatal synapses, enact essential functions that cannot be subsumed by cortex or any other parts of the motor system.

### New role for thalamus in learned motor sequence execution

Previously, DLS-projecting thalamic neurons have mainly been studied and discussed in terms of how they modulate signal flow and plasticity at corticostriatal synapses and how they contribute to attention and behavioral flexibility^52,64,158–160^. Electrophysiological recordings have shown that striatum-projecting thalamic neurons are active during sudden changes in behavioral context^62,63,160^, such as during contingency changes^161,162^ or self-initiation of movements^58,59^. Together, these results are consistent with thalamus providing a state and/or context-related signal which allows associations between the environment and appropriate movements and actions to be learned^58,160^.

Our results add to our understanding of the thalamostriatal pathway and its function by implicating DLS-projecting thalamic neurons in the control of learned motor sequences. However, much is left to sort out. Our silencing approach targeted the main DLS-projecting thalamic nuclei, i.e. the parafascicular nucleus and the rostral intralaminar and midline nuclei^49,51–54^. Because projections originating from different thalamic nuclei^25,52,163^ and even from different subpopulations within these nuclei^49^, can have distinct properties and functions^51,52,59,60,164,165^, it remains to be seen which of these projections play a role in motor sequence execution, and whether they have distinct functions. Furthermore, while the totality of our results points to plasticity at thalamostriatal synapses as an important mechanism for motor learning, this too must be conclusively established. Whether this is the main site of plasticity in DLS relevant for storing the memories of the behaviors we train, or whether there are contributions from corticostriatal plasticity at synapses formed by non-motor cortex neurons, remains to be investigated. Similarly, what role in execution of learned motor sequences, if any, the collaterals of DLS-projecting thalamic neurons play, must also be addressed.

This study probed the interactions between different brain areas within a distributed recurrent system (that includes thalamus, BG and motor cortex) to arrive at a mechanistic understanding of how learning and control algorithms are implemented. The underlying premise here is that complex processes can be broken down into component functions that can be attributed or localized to specific brain regions, pathways, and cell types. However, there can be no certainty that such mechanistic explanations are even possible in highly recurrent and distributed networks where functionality may be an emergent property of the system^166^. The striking dissociations we find, between the functions of different subregions of the striatum (DMS and DLS), and between its two main inputs from motor cortex and thalamus, can be seen as further evidence for modularity and separation of function in the mammalian motor system. While this may give some hope to those of us working towards a mechanistic understanding of motor learning and motor control, the path ahead remains perilous and long. Our hope is that the results presented here, and the insight that can be derived from them, will provide inspiration and guidance for future studies into the principles and mechanisms of how neural circuits underlie the acquisition and control of motor skills.

## Methods

### Animals

The care and experimental manipulation of all animals were reviewed and approved by the Harvard Institutional Animal Care and Use Committee. Experimental subjects were female Long Evans rats 3-10 months old at the start of training (n=68, Charles River). Because the behavioral effects of our circuit manipulations could not be pre-specified before the experiments, we chose sample sizes that would allow for identification of outliers and for validation of experimental reproducibility. Animals were excluded from experiments post-hoc if the lesions or infected areas were found to be outside of the intended target area or extended into additional brain structures (see Lesion section). The investigators were not blinded to allocation during experiments and outcome assessment, unless otherwise stated.

### Behavioral Training

Rats were trained in a lever-pressing task as previously described^35^. Water-restricted animals were rewarded with water for pressing a lever twice within performance-dependent boundaries around a prescribed interval between the presses (IPI = 700 ms). In addition, animals had to withhold pressing for 1.2 s after unsuccessful trials before initiating a new trial (inter-trial interval ITI). All animals were trained in a fully automated home-cage training system^74^. Manipulations were either performed in naïve animals before beginning of the training (Fig. 2) or after they had reached our learning criteria (mean IPI = 700 ms +/− 10%; CV of IPI distribution < 0.25; see Behavioral analysis below) and a median ITI >1.2 s, indicating that they had learned the task structure and stabilized their performance (Figs. 3–5).

### Lesion surgeries

Bilateral striatal lesions in naïve animals, targeting either the motor cortex-recipient part (DLS) or the non-MC input receiving part (DMS), were performed in two stages with a 10-day break between surgeries. Lesions were performed as previously described^35,167^. Briefly, animals were anesthetized with 2% isoflurane in carbogen and placed in a stereotactic frame. After incision of the skin along the midline and cleaning of the skull, Bregma was located and small craniotomies for injections were performed above the targeted brain areas. A thin glass pipette connected to a micro-injector (Nanoject II, Drummond) was lowered to the injection site and the excitotoxin quinolinic acid (0.09M in PBS (pH=7.3), Sigma-Aldrich) was injected in 4.9 nl increments to a total volume of 175 nl per injection site, at a speed of < 0.1 ul/min. After injection, the glass pipette was retracted by 100 um and remained there for at least 3 min before further retraction to allow for diffusion and to prevent backflow of the drug. After all injections were performed, the skin was sutured and animals received painkillers (Buprenorphine, Patterson Veterinary). Animals were allowed to recover for 10 days after the second surgery before being put into training.

To test for nonspecific effects of surgery and striatal injections on behavior, we performed 1-stage control surgeries according to the procedure described above, bilaterally injecting different non-toxic solutions, either fluorophore-coated latex microspheres (red excitation [exc.] = 530 nm, emission [em.] = 590 nm and green exc. = 460 nm, em. = 505 nm) referred to as retrobeads (Lumafluor)^168,169^ or Adeno-associated viruses (AAVs) for non-specific expression of GFP (Penn Vector Core), into DLS. This allowed for post-hoc evaluation of the targeting of our control injections.

Injection coordinates were (in mm, according to Paxinos^170^):

|  | DLS |  |  |  | DMS |  |  |  |
| --- | --- | --- | --- | --- | --- | --- | --- | --- |
| AP* | +0.7 | +0.7 | -0.3 | -0.3 | +1.2 | +1.2 | +0.2 | +0.2 |
| ML* | +/-3.6 | +/-3.6 | +/-4.0 | +/-4.0 | +/-1.9 | +/-1.9 | +/-1.9 | +/-1.9 |
| DV | -5.5 | -4.5 | -5 | -4 | -5.5 | -4.5 | -5.5 | -4.5 |

\* measured from bregma

### ZIP injections

To inhibit the enzyme PKMzeta and to reverse synaptic plasticity in different target regions, we injected the inhibitory peptide ZIP^98,99^ into either motor cortex, DLS or DMS. Animals were injected once they had reached our learning criteria (see Behavioral Training). We performed 1-stage injection surgeries according to the procedure described above. ZIP (10 mM in PBS, Tocris) was injected in 10 nl increments to a total volume of 500 nl per injection site^88,100^. To achieve a wider spread of the injected drug, injections were performed over a dorsal-ventral range by slowly moving the injection pipette dorsally while continuously injecting at evenly spaced intervals. Injection sites were verified post-hoc by locating co-injected fluorescent beads (data not shown). No animals had to be excluded based on the injection sites. Animals were put back into training after 5 days of recovery.

To estimate the spread of ZIP in the different brain areas, we injected a separate cohort of animals with ZIP-Biotin (10 mM in PBS, Tocris)^97^ which could be visualized post-hoc. We note that ZIP-Biotin has a higher molecular weight than uncoupled ZIP and is therefore expected to diffuse less far, providing only a lower bound for the actual spread of uncoupled ZIP in our experimental animals. Furthermore, the time course of the diffusion of uncoupled ZIP and the time of the biggest extent of its spread are unclear. We therefore determined the spread of ZIP-Biotin at two timepoints. Animals were perfused either 2h after ZIP injection (n=1 each for MC, DMS, DLS) or 24h after injection (n=1, DLS). We used either FITC-coupled avidin (1:100 in blocking solution (see Histology), Thermo Fisher Scientific) or fluorescently labeled anti-biotin antibodies (1:400 in blocking solution (see Histology), Jackson Immuno) for visualization of alternate slices of the individual brains (see Histology).

Injection coordinates were (in mm, according to Paxinos^170^):

|  | MC |  |  |  |  | DMS |  |  | DLS |  |  |  |
| --- | --- | --- | --- | --- | --- | --- | --- | --- | --- | --- | --- | --- |
| AP* | +1 | +1 | +1.5 | +2.25 | +3 | +1.5 | +1 | +0.2 | +1.75 | +1 | -0.25 | -0.7 |
| ML* | +/-2 | +/-4 | +/-2.75 | +/-2.5 | +/-2 | +/-1.9 | +/-1.9 | +/-1.9 | +/-3.2 | +/-4 | +/-4 | +/-4 |
| DV | -1.5 to -0.7 | -1.5 to -0.7 | -1.5 to -0.7 | -1.5 to -0.7 | -1.5 to -0.7 | -5 to -3.4 | -5.8 to -4 | -5.8 to -3.8 | -4.8 to -3.5 | -4.5 to -3.5 | -5 to -3.5 | -5 to -3.8 |

\* measured from bregma

### Projection-specific silencing of synaptic transmission

To silence neurons either in motor cortex or in thalamus which send axons to the DLS, we used a two-component viral strategy. We injected viruses, which can retrogradely infect neurons by entering axon terminals, into the DLS (either canine adeno virus (CAV)^79^ (IGMM, Montpellier) or retrograde AAV^80^ (Janelia, Addgene). These viruses drive expression of Cre recombinase in infected neurons. We further injected AAVs for Cre-dependent conditional expression of Tetanus Toxin Light Chain coupled to GFP (TeLC-GFP)^81–83^ (DNA construct shared by P. Wulff (University of Kiel), custom virus production by Harvard Medical School, Ocular Genomics Institute, Gene Transfer Core) into either motor cortex or thalamus. While the AAVs non-specifically infect neurons in the target area, TeLC is only expressed in neurons expressing Cre, i.e. neurons infected by the retrograde viruses. This strategy allowed us to target cortical or thalamic projection neurons which send an axon to the DLS. TeLC cleaves the synaptic protein VAMP2, thereby preventing the fusion of synaptic vesicles with the membrane and the release of neurotransmitters, effectively silencing synaptic transmission in infected neurons. Injections were performed as 1-stage surgeries, either in naïve (Fig. 2) or expert animals (Figs. 4, 5) and were given 5 days for recovery from surgery before (re-)start of training. Spread of the TeLC expression was determined post-hoc (see Histology).

To estimate the infection rate of our viral approach we co-injected retrobeads (Lumafluor) (see above: Lesion surgeries) into DLS in a subset of animals. Retrobeads are taken up by axonal terminals and transported retrogradely with high efficiency^168,169^, allowing us to use the number of retrobead-labeled neurons in cortex or thalamus as an estimate for the number of neurons projecting to DLS from these areas. We determined the efficiency of our viral approach by comparing the numbers of retrobead-labeled and GFP-expressing neurons at the injection sites in cortex or thalamus. We counted neurons in regions of interest in motor cortex or thalamus in 3 slices per animal (n=2 animals each for cortex and thalamus). Infection rates were similar between cortex and thalamus and reached about 50% at the centers of the injection sites.

Injection coordinates were (in mm, according to Paxinos^170^):

|  | MC |  |  |  |  | DLS |  |  |  |
| --- | --- | --- | --- | --- | --- | --- | --- | --- | --- |
| AP* | +1 | +1 | +1.5 | +2.25 | +3 | +1.75 | +1 | -0.25 | -0.7 |
| ML* | +/-2 | +/-4 | +/-2.75 | +/-2.5 | +/-2 | +/-3.2 | +/-4 | +/-4 | +/-4 |
| DV | -1.5 to -0.7 | -1.5 to -0.7 | -1.5 to -0.7 | -1.5 to -0.7 | -1.5 to -0.7 | -4.8 to -3.5 | -4.5 to -3.5 | -5 to -3.5 | -5 to -3.8 |

|  | Thalamus |  |  |  |  |
| --- | --- | --- | --- | --- | --- |
| AP* | -3.3 | -3.3 | -3.8 | -3.8 | -4.2 |
| ML* | +/-1.2 | +/-0.4 | +/-1.4 | +/-0.4 | +/-1.4 |
| DV | -5.8 to -4.3 | -6.2 to -5.8 | -6.2 to -4.3 | -6.1 to -5.9 | -6.1 to -4.8 |

\* measured from bregma

### Histology

At the end of the experiment, animals were euthanized (100 mg/kg ketamine and 10 mg/kg xylazine, Patterson Veterinary), perfused with 4% paraformaldehyde (PFA), and their brains were harvested for histology to confirm lesion size, injection sites and drug/ viral spread. The brains were sectioned into 80 μm slices and stained in different ways. To determine lesion location and size, slices were stained with Cresyl Violet following standard procedures. In a subset of animals immunofluorescence stainings were performed instead of Cresyl Violet stainings. After slicing, sections were blocked for 1h at room temperature in blocking solution (1% BSA, 0.3% TritonX), stained overnight at 4C with primary antibodies for NeuN (to stain for neuronal cell bodies; 1:500 in blocking solution; Millipore MAB377) and GFAP (to stain for glia cells; 1:500 in blocking solution; Sigma) and subsequently with appropriate fluorescently-coupled secondary antibodies (1:1000 in blocking solution) for 2h at room temperature. The same staining protocol was used to visualize TeLC-GFP, using antibodies for NeuN and GFP (1:1000 in blocking solution, Life Technologies) or to visualize ZIP-Biotin, using antibodies for Biotin (1:400 in blocking solution, Jackson Immuno). In alternate slices, ZIP-Biotin was visualized by incubation with fluorescein-coupled Avidin (1:100 in blocking solution, Thermo Fisher) over night. Images of whole brain slices were acquired at 10x magnification with either a VS210 Whole Slide Scanner (Olympus) or an Axioscan Slide Scanner (Zeiss).

### Quantification of lesion size, viral infection and ZIP spread

To determine the extent and location of striatal lesions, we analyzed several sections (4-6) spanning the anterior-posterior extent of the striatum, allowing for an estimate of the overall lesion size. Lesion boundaries were determined throughout the striatum and adjacent areas, blind to the animals’ identity and performance. Boundaries were marked manually based on differences in cell morphology and density (loss of larger neuronal somata and accumulation of smaller glial cells). The extent of the striatum was determined based on the Paxinos Rat Brain Atlas, using anatomical landmarks (external capsule, ventricle) and cell morphology and density. Additionally, we marked the GPe in posterior sections, since mistargeted injections may lead to its partial lesioning, disrupting the output both of the DLS and DMS.

In addition to the overall lesion size, we also determined the lesioned fractions of the DLS/DMS. Since the DLS and DMS are not clearly defined, we made use of their differential input patterns from MC and PFC, respectively, which we had previously determined using viral anterograde labeling^11^. We used these identified boundaries of the DLS and DMS to determine the lesioned fractions in the experimental animals. We pre-defined a threshold based on our previous observations ^11^ of at least 50% loss of the targeted region, less than 10% loss of the non-targeted part of the striatum and no lesions in the GPe for inclusion of experimental animals in our analysis. Based on this threshold, no animals had to be excluded.

To determine the spread of TeLC expression in our silencing experiments, we determined the affected areas in motor cortex and thalamus, respectively. We manually labeled the extent of the infections based on the presence of GFP-expressing somata in the respective regions. Animals with no discernable expression of GFP in cortex or in thalamus were excluded from behavioral analysis (n=2 and n=1, respectively). We further used co-injected fluorescent beads to verify the injection sites of the retrograde viruses in the DLS.

To determine a lower boundary for the spread of ZIP injections in the different brain areas, we manually labeled the extent of fluorescent labeling around the injection sites in the cohort of animals injected with ZIP-Biotin. In experimental animals injected with non-labeled ZIP, we used co-injections of retro-beads to verify the injection sites in the respective target areas. Based on this, no animals had to be excluded.

### Kinematic Tracking

To determine the movement trajectories of the forelimbs of animals performing our task, we made use of recently developed machine learning approaches, using deep neuronal networks to determine the position of specific body parts in individual video frames^107^,108.

Videos of animals performing the task were acquired at 30Hz and saved as snippets ranging from 1s before the first lever press in a trial to 2s after the last lever press in the trial. We extracted about 1000 frames randomly selected throughout the duration of the trials, balanced across pre- and post-manipulation conditions and manually labeled the position of the forelimbs in each frame, using custom-written Matlab code. This data was used to train individual neural networks for each animal.

We trained ResNet-50 networks that were pretrained on ImageNet, using the DeeperCut implementation in TensorFlow (https://github.com/eldar/pose-tensorflow^107^. Training was performed using default parameters (1 million training iterations, 3 color channels, with pairwise terms, without intermediate supervision). Data augmentation was performed during training by rescaling images from a range of 85% to 115%.

The trained neural network was then used to predict the position of the forelimbs in all trials across conditions, on a frame-by-frame basis. The position of a forelimb in a frame is given by the peak of the network’s output score-map. Frames in which the forelimb was occluded were identified as having a low peak score. For both the training and the subsequent predictions we used GPUs in the Harvard Research Computing cluster.

Because the two forelimbs could often be confused for each other in the neural network’s predictions from a single frame, we took advantage of correlations across time to constrain the predictions. For each forelimb, the predicted score-maps for all frames in a single trial video were passed through a Kalman filter using the Python toolbox *filterpy*. Specifically, a constant-acceleration Kalman smoother was used which assumes that the forelimb on adjacent frames will have the same acceleration (zero jerk) plus a small noise term. Only frames with a weak neural-network prediction score were adjusted by the Kalman filter; otherwise the original neural-network prediction was used as the forelimb position.

The tracking accuracy was validated post-hoc by visual inspection of at least 50 predicted trajectories per animals. Initial training with lower frame numbers often led to inaccurate tracking results. After settling on a 1000 training frames, none of the trained networks was discarded.

Missing frames in the trajectories, e.g. due to temporary occlusions of the forelimbs, were linearly interpolated for a maximum of 5 consecutive frames. If occlusions lasted longer, the trajectories were discarded. In a subset of animals, the quality of the recorded videos was not sufficient for high-quality tracking of the forelimbs, due to inappropriate lighting conditions or due to occlusions of the forelimbs over long durations of the trials and we had to discard the trajectories (n=4).

### Behavioral Data Analysis

#### Performance Metrics

Performance metrics were determined based on the timing of lever presses in our task. The inter-press interval (IPI) was determined as the time between the first and second press in a trial, the inter-trial interval (ITI) as the time between the last press in an unsuccessful trial and the next occurring lever press. The CV during learning (Figs. 2, S2) was calculated across 100 trials and the moving average was low pass-filtered with a 300-trial boxcar filter. For the manipulations in expert animals (Figs. 3, 4, S4, S6) a smaller moving window (25 trials) and boxcar filter (50 trials) were used. The fraction of trials close to the target IPI was calculated using the same windows and filters. Trials were labeled as close to the target if they were in the IPI range of 700 ms +/− 20%.

### Criterion Performance

We considered animals as having successfully learned the task and as having reached criterion performance if the CV was below 0.25 and the mean of the IPI distribution was in the range of 700 ms +/− 10% for a 3000 trial sliding window.

### JS Divergence

As a measure for the dissimilarity of the IPI and ITI distributions in individual animals, we calculated the Jensen-Shannon (JS) Divergence of the distributions. The JS Divergence is a symmetric derivative of the Kullback-Leibler divergence (KLD). We calculated the JS Divergence (JSD) as:

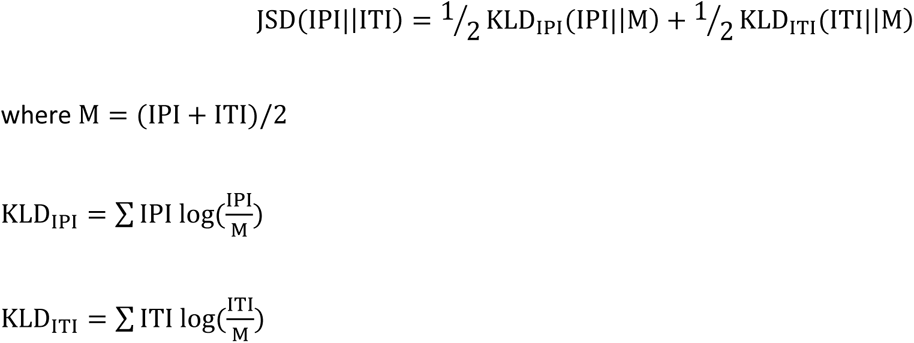

### Movement trajectory analysis

We compared the trajectories of both forelimbs of all tracked animals before and after Th-DLS silencing (Fig. 5). We focused on the position of the forelimbs in the vertical dimension, in which the movements in our task are more pronounced than in the horizontal dimension. To be able to compare the stereotypy of the trajectories for the learned motor sequences, we sub-selected trials which were successful and rewarded, and which occurred after unrewarded trials. This allowed us to compare trials with the same start and end positions. This is necessary, since animals move down to, and back up from, a reward port underneath the lever after successful trials^35^. Some animals learn more than one stereotyped motor sequence as solutions to our task. These solutions, to which we refer as ‘modes’, occur at different frequencies, usually with one mode being predominant, but all become stereotyped and have similar properties. For our analysis we sub-selected only trials from the most common mode, so that trajectories are comparable. We determined the modes by hierarchical clustering of the time-warped forelimb trajectories^11^. We plotted the average of the selected trajectories for before and after the manipulation, calculated the SEM (Fig. 5A) and plotted a projection of all selected trials (Fig. 5B). To calculate the correlations between the individual trials, we linearly warped the trajectories to the same duration by interpolating between the lever presses. Since the lever presses themselves have stereotyped trajectories, largely independent of the trial duration, we interpolated only the trajectories from 100 ms after the first to 100 ms before the second lever press to preserve the shape of the presses. From these warped trajectories we calculated trial-to-trial correlations separately for both forelimbs and averaged the correlations for each trial (Fig. 5C). These correlations were averaged for the individual conditions within animals and those means were averaged across animals and plotted with the SEM (Fig. 5D).

To compare the trajectories across animals, we linearly warped all trajectories and normalized their amplitude to their individual maximum amplitude (Fig. 5E,F). To calculate the correlations across animals, we first calculated the average pair-wise correlations across all trials within individual animals, and then averaged these across the individual animals (Fig. 5G,H).

We separately compared the lever press movements, defined as the trajectory in the range of +/− 150 ms around a detected lever press (Fig. 5I,J). We normalize the lever press trajectories to their individual maximum amplitude and plotted their overlay (Fig. 5I). To compare the lever presses before and after the manipulation to the presses early in training, we additionally sub-selected trials as described above from the first 2000 trials of training. As above, we calculated the average pairwise correlations for all lever presses in all trials of all animals across the conditions (early, pre- and post-silencing) and averaged them first by lever press (i.e. animal 1 press 1, animal 1 press 2, etc.) and then by condition (Fig. 5J).

### Statistical analysis

All statistics on data pooled across animals is reported in the figures as mean +/− SEM. Multiple comparison tests were used where justified. Statistical tests for specific experiments were performed as described below.

*Fig. 2D.* Reaching of the learning criterion was compared between control and DMS-lesioned animals. A two-tailed unpaired t-test revealed no significant differences in reaching the learning criterion between the control and DMS-lesioned animals (n=6/4; P = 0.1179).

*Fig. 2F.* The JS divergences between the IPI and ITI were compared between early and late in learning. A two-tailed paired t-test revealed significant differences for the control (n=6; P =0.0088) and DMS-lesioned groups (n=4; P =0.005), but neither for the DLS-lesioned (n=5; P =0.7537), nor the MC-DLS silencing group (n=9; P =0.4087).

*Fig. 3E.* The JS divergences between the IPI and ITI were compared between pre- and post-ZIP for the MC and DMS groups, using a two-tailed paired t-test revealing no significant differences (MC: n=7; P = 0.92; DMS: n=5; P =0.06). For the DLS group the JS divergences were compared across 4 conditions (early, pre-ZIP, post-ZIP, late after ZIP). A repeated measures ANOVA revealed significant differences between time points (F(3,15)=7.86, P=0.038). Post-hoc comparisons using the Bonferroni test showed significant differences between early and pre-ZIP (P=0.008) and between pre- and post-ZIP (P=0.012).

*Fig. 4E.* The JS divergences between the IPI and ITI were compared between pre- and post-silencing for the Control and MC groups, using a two-tailed paired t-test revealing no significant differences. For the thalamus group the JS divergences were compared across 4 conditions (early, pre-silencing, post-silencing, late after silencing). A repeated measures ANOVA revealed significant differences between time points (F(3,24)=13.37, P=0.0064). Post-hoc comparisons using the Bonferroni test showed significant differences between early and pre-silencing (P<0.001), between pre- and post-silencing (P<0.001) and between pre-silencing and late (P=0.001).

*Fig. 5D.* The within trial-to-trial pairwise correlations per animal were compared for pre- and post-silencing and between pre- and post-silencing. A repeated measures ANOVA revealed significant differences between time points (F(2,6)=8.17, P=0.0194). Post-hoc comparisons using the Bonferroni test showed significant differences between pre and pre-post (P=0.042) and between post and pre-post (P=0.035).

*Fig. 5H.* The pairwise correlations for the forelimb trajectories were calculated between all trials across animals. These correlations were averaged for individual animals and all correlations per condition were averaged. A repeated measures ANOVA revealed significant differences between time points (F(2,6)=18.40, P=0.0028). Post-hoc comparisons using the Bonferroni test showed significant differences between pre and post (P=0.004) and between post and pre-post (P=0.009).

*Fig. 5J.* The pairwise correlations for the lever press trajectories were calculated between all trials across animals. These correlations were averaged for individual animals and all correlations per condition were averaged. A repeated measures ANOVA revealed significant differences between time points (F(5,35)=17.13, P<0.0000). Post-hoc comparisons using the Bonferroni test showed significant differences between early and pre (P=0.003), early and early-pre (P<0.001), early and pre-post (P=0.004), pre and post (P<0.001), pre and early-post (P=0.021), post and earl-pre (P<0.001), post and pre-post (P<0.001), early-pre and early-post (P<0.001) and early-post and pre-pose (P=0.024).

*Fig. S2B.* The IPI was compared between early and late learning for different performance metrics. Two-tailed paired t-tests revealed significant differences for control (n=6; P=0.0147) and DMS (n=4; P=0.0438) animals. For the CV two-tailed paired t-tests revealed significant differences for control (n=6; P<0.001) and DMS (n=4; P=0.042) animals. For the Precision (trials close to target) two-tailed paired t-tests revealed significant differences for control (n=6; P<0.001) and DMS (n=4; P=0.0109) animals. For the ITI two-tailed paired t-tests revealed significant differences for control (n=6; P<0.001) and DMS (n=4; P=0.0032) animals.

*Fig. S4B.* For the comparison of performance metrics before and after ZIP for the MC and DMS group, two-tailed paired t-tests were used and did not yield any significant difference for any of the comparisons. For the DLS group 4 time points were compared (early, pre-ZIP, post-ZIP, late after ZIP). For the IPI a repeated measures ANOVA revealed significant differences between time points (F(3,18)=10.62, P=0.0003). Post-hoc comparisons using the Bonferroni test showed significant differences between early and pre (P<0.001) and early and late (P=0.001). For the CV a repeated measures ANOVA revealed significant differences between time points (F(3,18)=24.04, P<0.0000). Post-hoc comparisons using the Bonferroni test showed significant differences between early and pre (P<0.001), early and post (P=0.010), early and late (P<0.001) and pre and post (P=0.003). For the Precision a repeated measures ANOVA revealed significant differences between time points (F(3,18)=34.3, P<0.0000). Post-hoc comparisons using the Bonferroni test showed significant differences between early and pre (P<0.001), early and late (P<0.001), pre and post (P<0.001) and post and late (P=0.002). For the ITI a repeated measures ANOVA revealed significant differences between time points (F(3,18)=9.6, P=0.0005). Post-hoc comparisons using the Bonferroni test showed significant differences between early and pre (P=0.003), early and late (P=0.008), pre and post (P=0.009) and post and late (P=0.023).

*Fig. S6B.* For the comparison of performance metrics before and after silencing for the Control and MC group, two-tailed paired t-tests were used and did not yield any significant difference for any of the comparisons. For the thalamus group 4 time points were compared (early, pre-silencing, post-silencing, late after silencing). For the IPI a repeated measures ANOVA revealed significant differences between time points (F(3,21)=4.75, P=0.0111). Post-hoc comparisons using the Bonferroni test showed significant differences between early and pre (P=0.014). For the CV a repeated measures ANOVA revealed significant differences between time points (F(3,21)=13.73, P<0.0000). Post-hoc comparisons using the Bonferroni test showed significant differences between early and pre (P<0.001), pre and post (P<0.001) and pre and late (P=0.006). For the Precision a repeated measures ANOVA revealed significant differences between time points (F(3,21)=16.38, P<0.0000). Post-hoc comparisons using the Bonferroni test showed significant differences between early and pre (P<0.001), pre and post (P<0.001) and pre and late (P<0.001). For the ITI a repeated measures ANOVA revealed significant differences between time points (F(3,21)=19.17, P<0.0000). Post-hoc comparisons using the Bonferroni test showed significant differences between early and pre (P<0.001), pre and post (P<0.001) and pre and late (P<0.001).

---

**Supplementary Figure 1:**
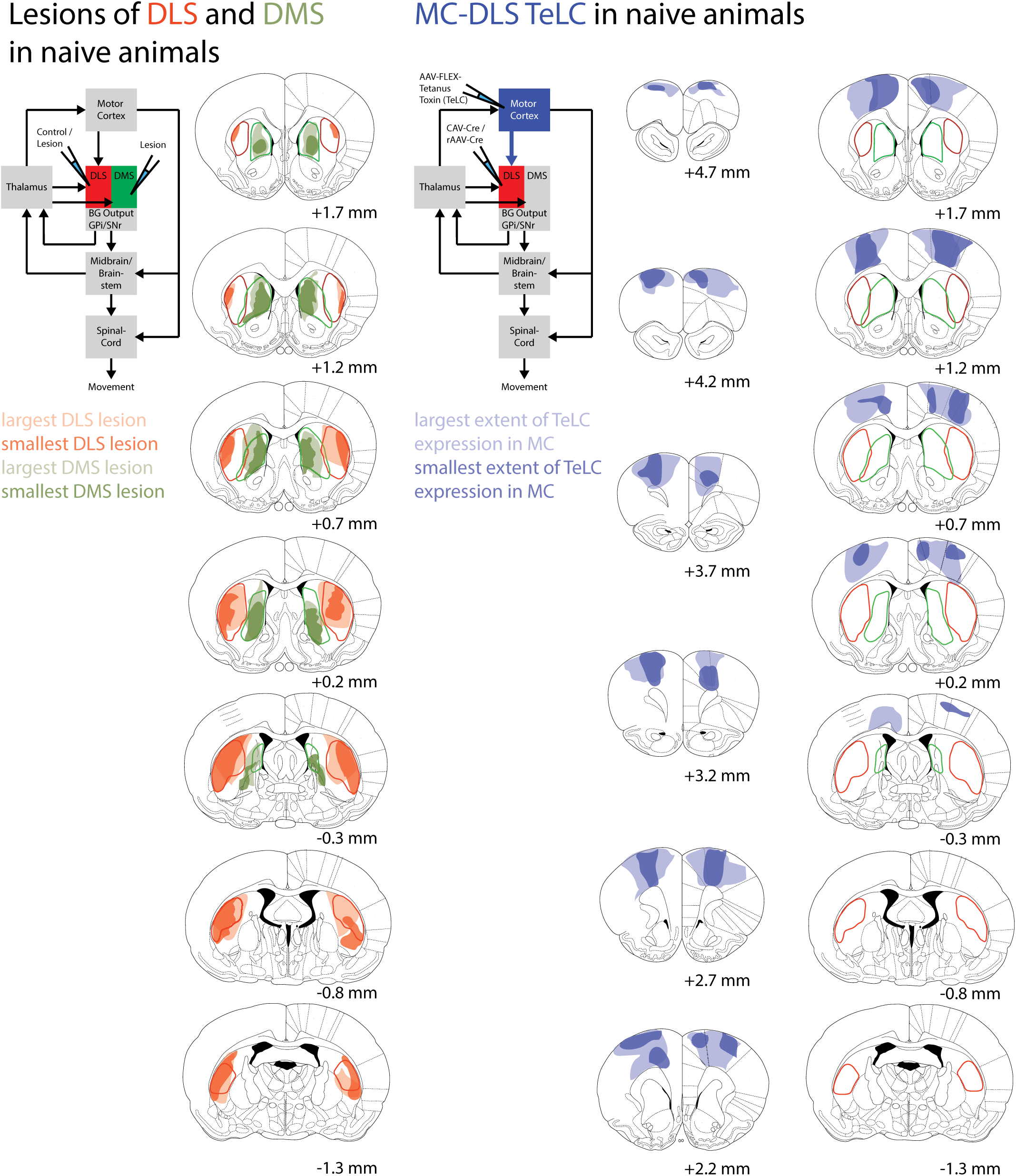
Extent of lesions of the DLS and DMS and of TeLC expression in motor cortex. *Left*: Red/green: DLS/DMS lesions. Light and dark colors indicate the extent of the largest and smallest lesion. *Right*: Light and dark blue: largest and smallest extent of TeLC expression. Red and green outlines mark the extent of motor and prefrontal cortex projections to the striatum as previously identified by viral anterograde labeling of projection fibers^11^.

**Supplementary Figure 2:**
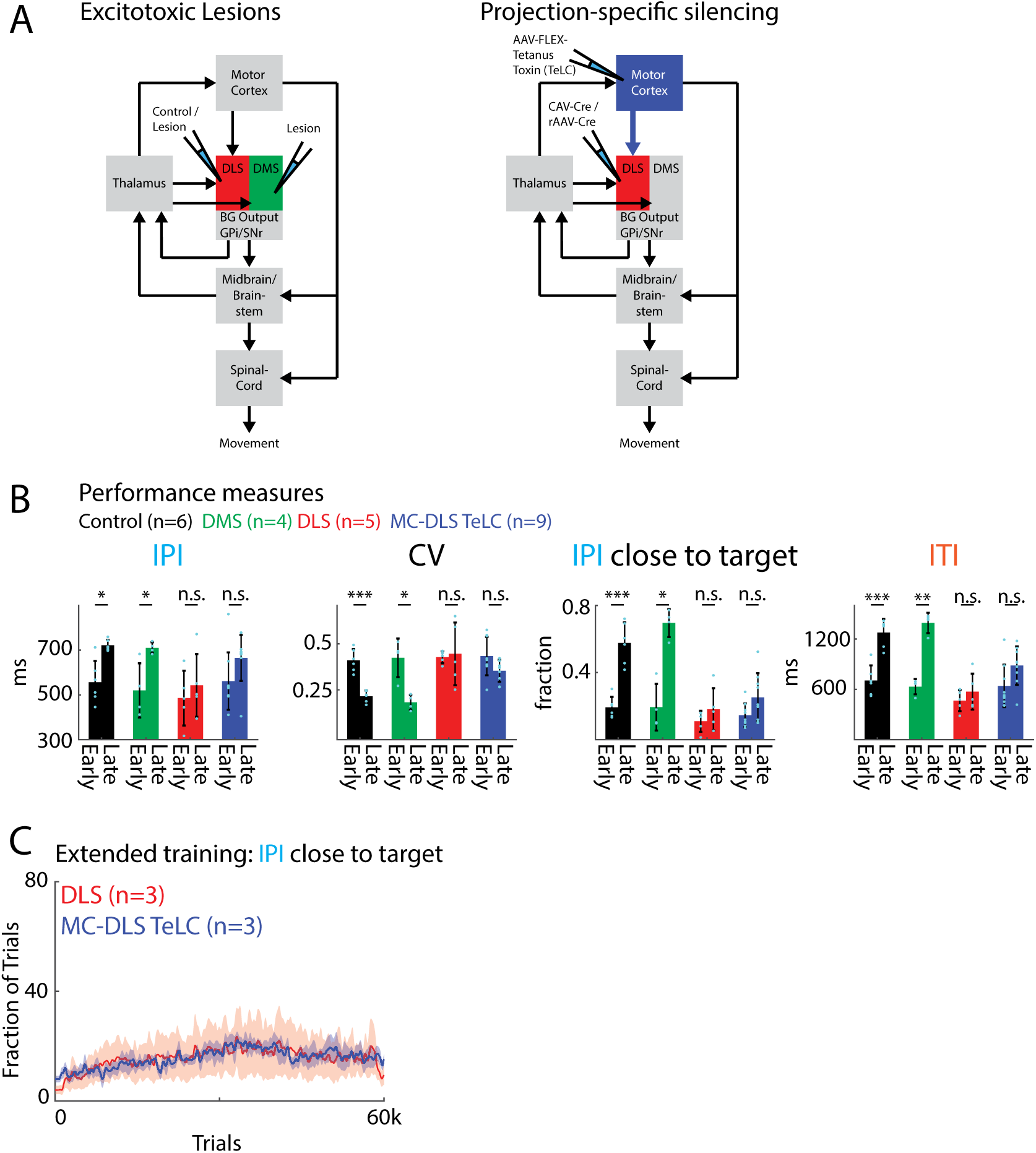
**A)** Experimental schemes for *Left*: excitotoxic lesions and *Right*: projection-specific silencing (see Fig. 2) **B)** Comparisons of performance measures between different manipulations early and late in training (as in Fig. 2), show an improvement in performance for control and DMS animals, but not for DLS and MC-DLS TeLC animals. IPI: inter-press interval, CV of IPI: Coefficient of Variation of the IPI, IPI close to target: Fraction of trials close to target IPI (700ms +/− 20%), ITI: inter-trial interval. Early: First 2000 trials in training, Late: trials 30,000 to 32,000 in training. Bars represent means across animals and dots represent means within individual animals. Error bars represent standard error of the mean (SEM). **C)** Population results for extended training in animals which underwent either DLS lesions or MC-DLS silencing. Even after 60,000 trials animals’ performance did not improve. Shown is the fraction of trials with IPI close to the target (700 ms +/− 20%). *P < 0.05, **P < 0.01, ***P < 0.001.

**Supplementary Figure 3:**
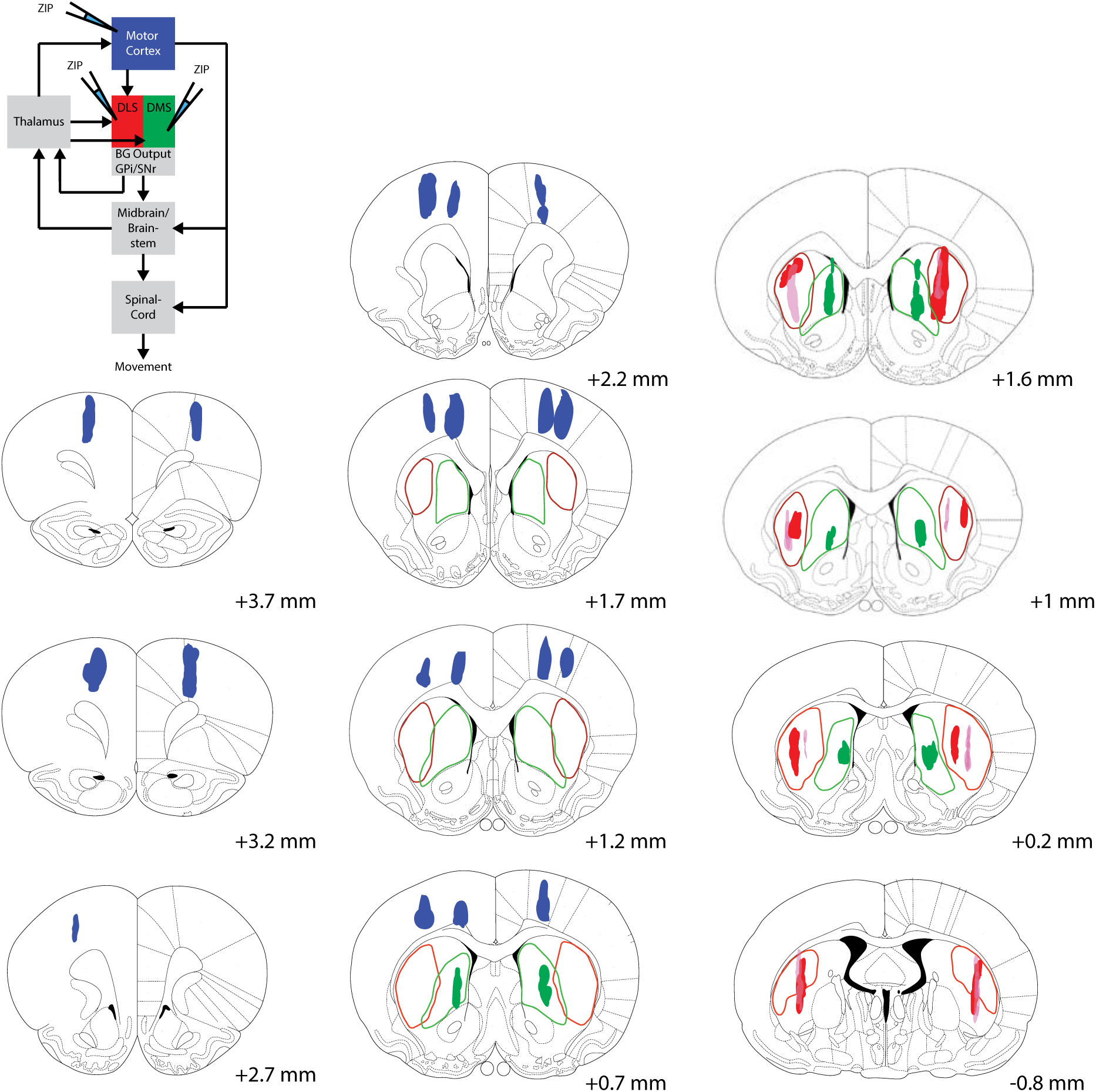
Spread of ZIP injections into motor cortex, DLS or DMS. Using biotinylated ZIP and fluorescently labeled avidin, we determined a lower bound for the spread of non-labeled ZIP (with lower molecular weight) in the different target areas. Blue: MC, Green: DMS, Red: DLS – dark and light colors indicate injections in different animals. Colored lines indicate the outlines of the DLS and DMS as previously determined by viral labeling of axons from motor cortex and prefrontal cortex, respectively^11^.

**Supplementary Figure 4:**
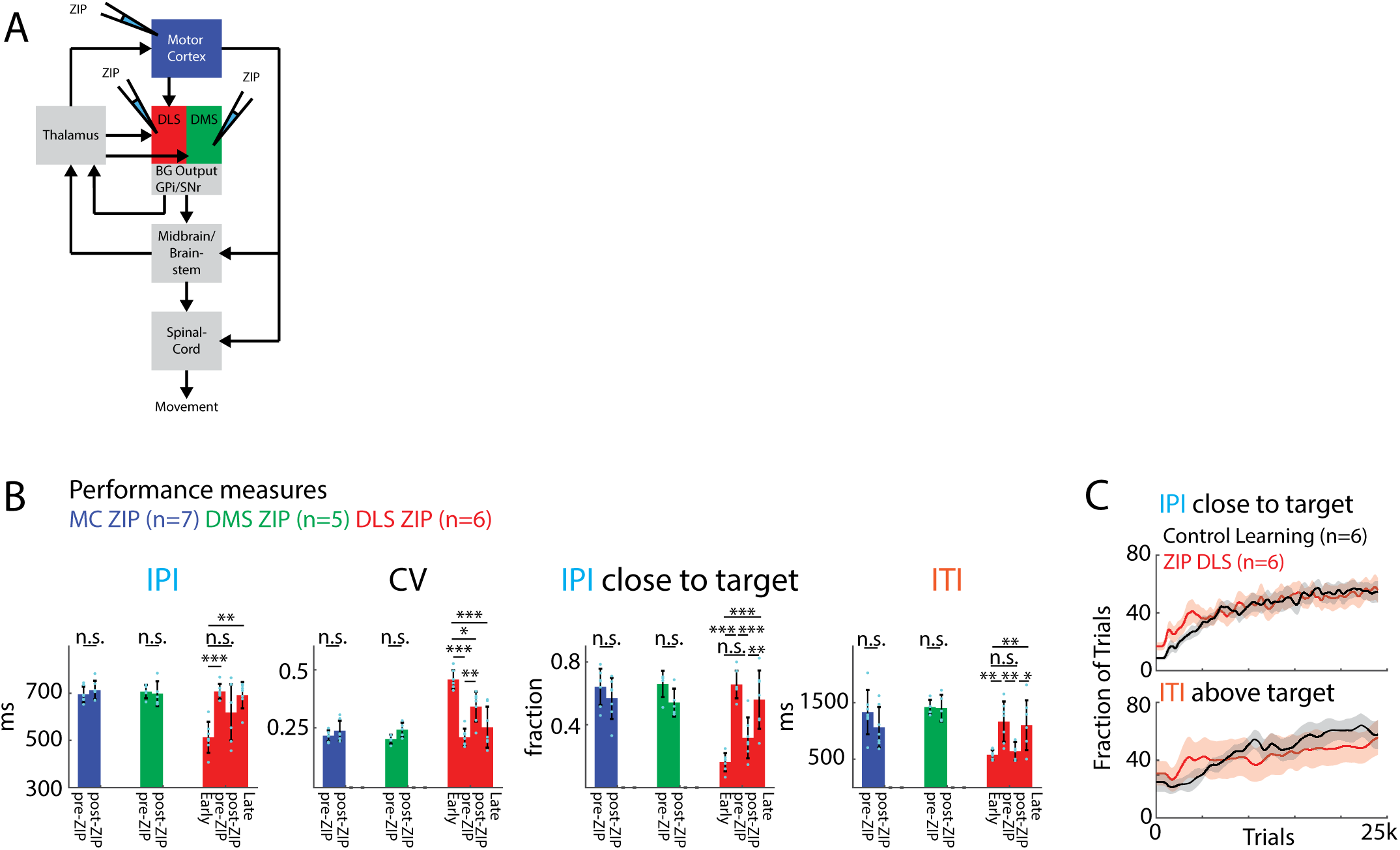
**A)** Experimental scheme for ZIP injections in motor cortex, DMS or DLS (see Fig. 3). **B)** Comparison of performance measures at different stages before and after ZIP injections, showing a reduced performance after DLS-ZIP to levels observed early in training. IPI: inter-press interval, CV of IPI: Coefficient of Variation of the IPI, IPI close to target: Fraction of trials close to target IPI (700 ms +/− 20%), ITI: inter-trial interval. Early: First 2000 trials in training, pre-ZIP: last 2000 trials before ZIP, post-ZIP: first 2000 trials after ZIP, Late: trials 10,000 to 12,000 after ZIP. Bars represent means across animals and dots represent means within individual animals. Error bars represent standard error of the mean (SEM). **C)** Development of the IPI and ITI after ZIP DLS injections, compared to the performance of control animals (compare Fig. 2) over the course of re-learning and initial learning, respectively. Both the development of the IPI and ITI after DLS ZIP resemble normal learning. *P < 0.05, **P < 0.01, ***P < 0.001.

**Supplementary Figure 5:**
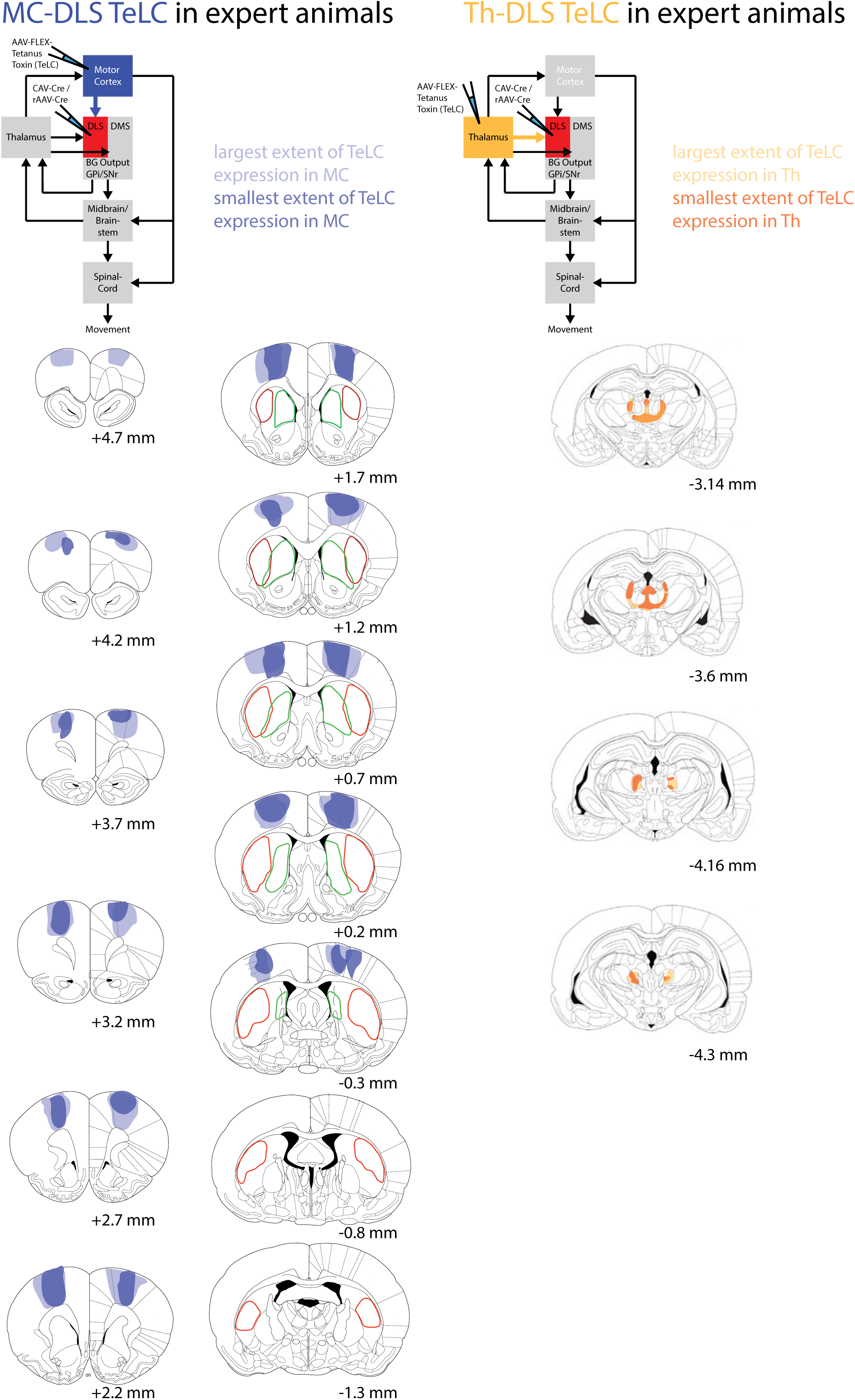
Extent of expression of TeLC in motor cortex or thalamus after projection-specific silencing. *Left*: Light blue: largest extent of expression of TeLC in motor cortex, Dark blue: smallest extent of expression. *Right*: Light yellow: largest extent of expression of TeLC in thalamus, Dark yellow: smallest extent of expression.

**Supplementary Figure 6:**
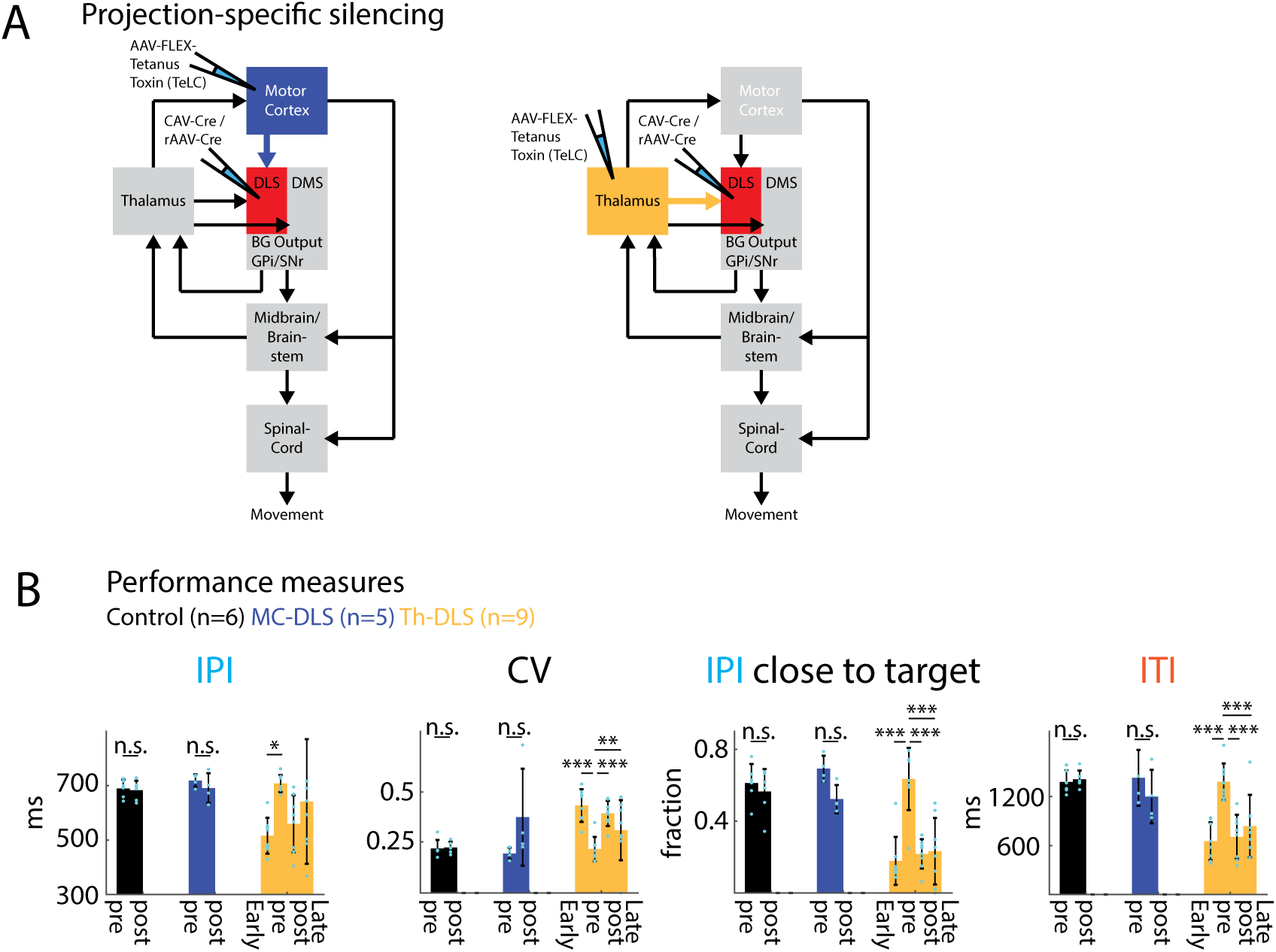
**A)** Experimental scheme for projection-specific silencing in motor cortex or thalamus (see Fig. 4) in expert animals. **B)** Comparison of performance measures at different stages before and after viral injections, showing a reduction of performance after Th-DLS silencing to levels observed early in training. IPI: inter-press interval, CV of IPI: Coefficient of Variation of the IPI, IPI close to target: Fraction of trials close to target IPI (700 ms +/− 20%), ITI: inter-trial interval. Early: First 2000 trials in training, pre-silencing: last 2000 trials before silencing, post-silencing: first 2000 trials after silencing, Late: trials 10,000 to 12,000 after silencing. Bars represent means across animals and dots represent means within individual animals. Error bars represent standard error of the mean (SEM). *P < 0.05, **P < 0.01, ***P < 0.001.

## Acknowledgements

We thank Ashesh Dhawale for advice on data analysis and Keven Laboy Juarez and Tobias Messmer for technical support. We further thank Sean Escola, James Murray and members of the Ölveczky lab for advice on data analysis and for discussions and comments on the manuscript. We thank Alexander Mathis and Mackenzie Mathis for introducing us to and initial help with kinematic tracking. We thank Steve Turney and the Harvard Center for Biological Imaging for infrastructure and support. We also thank Peer Wulff (University of Kiel) for sharing the TeLC construct and Alla Karpova (Janelia) and Adam Hantman (Janelia) for sharing the rAAV. This work was supported by NIH grants R01-NS099323-01 and R01NS105349 to B.P.Ö. and by EMBO and HFSP postdoctoral fellowships to S.B.E.W.

## Author Contributions

S.B.E.W. and B.P.Ö. designed the study. S.B.E.W. conducted all experiments and analyzed the data with help from B.P.Ö. R.K. performed pilot lesion experiments. S.B.E.W. and B.P.Ö. wrote the manuscript.

